# Evolution of m6A related genes in insects and function of METTL3 in embryonic development of silkworm

**DOI:** 10.1101/2022.08.29.505623

**Authors:** Shuai-Qi Liu, Shun-Ze Jia, Ying-Hui Li, Kai-Wen Hu, Jian-Guo Tao, Yi-Cheng Lu, Yu-Song Xu, Hua-Bing Wang

## Abstract

N6-methyladenosine (m6A), the most prevalent modification in eukaryotic RNAs, plays a key role in biological processes. However, the evolutionary relationships and function of m6A related genes in insects are still largely unknown. Here, we analyzed the phylogenetics of m6A related genes among 207 insects and find that m6A related genes are evolutionarily conserved in insects. We apply the lepidopteran model insect *Bombyx. mori* as a study system. Subcellular localization experiments in BmN cells confirmed that YTHDF3 localizes in the cytoplasm, METTL3, METTL14 and YTHDC localize in the nucleus, while FL2D localizes in both nucleus and cytoplasm. We also detected expression pattern of m6A related genes during the embryonic development of *B. mori* and found that m6A related genes expression pattern is temporally specific. To elucidate the function of METTL3 during embryonic stage, RNA sequencing was performed to measure differential expressions of mRNA from silkworm eggs after *METTL3* knockdown, and METTL3-overexpressing BmN cells. The global transcriptional pattern showed that *METTL3* knockdown affected multiple cellular processes, including oxidoreductase activity, transcription regulator activity, cation binding and fundamental many metabolism processes, such as carbon metabolism, purine metabolism, biosynthesis of amino acids, and the citrate cycle, were affected. In addition, *METTL3* knockdown significantly affected Wnt and Toll/Imd pathways in embryonic stage. The results suggest that METTL3 plays an important role during the embryonic development of *B. mori*. This study deepens our understanding of the function of m6A related genes in insects.

## Introduction

As the most common types of eukaryotes in RNA modification (Alarcón et al., 2015; Chandola et al., 2015; Holoch and Moazed, 2015), N6-methyladenosine (m6A) is the most abundant and reversible modification of various types of RNA, such as mRNA, transport RNA (tRNA), ribosomal RNA (rRNA)(Fustin et al., 2013; Holoch and Moazed, 2015). m6A is dynamically regulated by methyltransferase complex, demethylases, and m6A-binding proteins (Frye et al., 2018; Yang et al., 2018). m6A coordinates many steps of RNA metabolism ranging from splicing, RNA processing, RNA transport, mRNA translation, and RNA degradation (Claudio R. Alarcón et al., 2015; Chen et al., 2018; Lin et al., 2019; Wang et al., 2015, 2019).

m6A pathway has been shown to play a role in neuronal functions and sex determination in *Drosophila melanogaster*. m6A is required for female-specific alternative splicing of *Sxl*, which determines female physiognomy, but also translationally represses male-specific lethal 2 (*msl-2*) to prevent dosage compensation in females(Haussmann et al., 2016). Physiological METTL3/YTHDF function is explicitly required within mushroom body neurons to mediate normal conditioned odor memory during aging. The m6A pathway was found to influence the expression of *P450* gene-*CYP4C64*, which in turn influences insecticide resistance in whitefly (Yang et al., 2021). A point mutation (A-206T) in the 5’UTR of *CYP4C64* is observed at high frequency in thiamethoxam-resistant *Bemisia tabaci* strains and the m6A modification at this position increases expression of *CYP4C64* and resistance.

In *B.mori*, m6A affects cell cycle and the arrangement and separation of chromosomes in BmN cells(Li et al., 2019). In addition, m6A related genes expression and m6A content were higher in diapause-destinated compared to nondiapause-destined strains, suggesting that m6A modification may provide significant epigenetic regulation of diapause-related genes in the silkworm (Jiang et al., 2019). After the relative genes of the m6A pathway, such as *METTL14, METTL3*, and *YTHDF3* knockdown, the BmN cells have been found to have a low resistance to BmNPV infection, which indicated that m6A modification might be a novel epigenetic mechanism that regulates BmNPV infection (Zhang et al., 2020). However, physiological functions of m6A in insect embryonic development remain unexplored.

In this study, we analyzed the evolutionary relationship of m6A related genes in insects. We investigated expression patterns of m6A related genes and subcellular localization of m6A related proteins in the lepidopteran model organism *B. mori*. To investigate the role of METTL3 in *B. mori* embryonic development, *METTL3* was knockdown at early embryonic stages of the silkworm by siRNA, and RNA-Sequencing (RNA-Seq) analysis was performed. These data showed that MAPK, Wnt, apoptosis, and Toll/IMD signaling pathways are downstream target of METTL3 in embryonic development of *B. mori*. By overexpressing METTL3 in BmN cells and RNA-Seq analysis, we further confirmed that METTL3 plays an important role in regulating organelle function and purine metabolism. Our research provides novel insight into evolutionary relationship of m6A methylation and biological function of METTL3 in insects.

## Methods and materials

### Genome-wide identification of m6A related genes

Two procedures were used to identify potential m6A related genes. The Hidden Markov Model (HMM) profile for m6A related proteins was downloaded from Pfam Database (http://pfam.xfam.org/) and was used as a query to scan the *B. mori* genome using HMMER software (version 3.1b2; http://hmmer.org/) with a default E-value. Other potential homologous m6A related protein sequences were collected using the Blast program (blastp) in the National Center for Biotechnology Information database (NCBI, https://www.ncbi.nlm.nih.gov/protein) using *B. mori* m6A related proteins as search queries (E-value < 1e^-5^). Subsequently, the potential m6A related proteins were further verified for the presence of the m6A related domain by simple modular architecture research tool (SMART, available online: http://smart.embl-heidelberg.de/) (Ponting et al., 1999) and conserved domain database (CDD, available online: http://www.ncbi.nlm.nih.gov/Structure/cdd/wrpsb.cgi)(Marchler-Bauer et al., 2012).

### Multiple sequence alignment and phylogenetic reconstructions of m6A related proteins

The phylogenetic relationships between m6A related proteins from *B. mori* and other insects were determined using the GUIDANCE 2.0 Server (http://guidance.tau.ac.il/ver2/) for multiple sequence alignment of full-length amino acid sequences. The best fit amino acid replacement model (LG+I+G), determined with ProtTest, was used to carry out phylogenetic analyses using the maximum-likelihood (ML) method in the MEGA 11.0.1 software with the rapid bootstrap algorithm (1 000 replicates)(Tamura et al., 2021). Phylogenetic trees were visualized using the Interactive Tree of Life (iToL, version 5.5.1, http://itol.embl.de)(Letunic and Bork, 2019). GenBank accession numbers of sequences used in this study are listed in Table S2-S6.

### RNA extraction and quantitative reverse transcription-PCR (qRT-PCR)

Total RNA from silkworm eggs, was isolated using RNAiso Plus (TaKaRa, Japan). PrimerScript RT reagent kit with gDNA Eraser (TaKaRa, Japan) was employed to reverse transcribe the total RNA (1 μg) according to the manufacturer’s instructions. To analyze gene expression, quantitative real-time PCR (qPCR) was performed as described previously. Specific primers for qRT-PCR analysis listed in TableS1 were designed using Primer 5 software. Expression levels for each sample were calculated from three biological replicates, each with three technical replicates. The expression of ribosomal protein 49 (*rpl49*) was used as a control to normalize gene expression. Student’s t-test was used for evaluating the statistical significance.

### Protein purification and determination of recombinant BmMETTL3(204-329)

CDS sequence coding protein BmMETTL3(204-329) was cloned into the Pcold vector (TaKaRa, China). Primers were listed in Table S1. *Escherichia coli* carrying BmMETTL3(204-329) were grown at 37°C in Luria-Bertani medium. After 18 h of incubation at 18°C and 160 rpm. The harvested bacteria were immediately centrifuged at 6000 rpm 4 °C for 10 min. The supernatant was collected after centrifugation. Next, the pellets of bacteria were resuspended in ice-cold lysis buffer containing protease inhibitors. The cells were lysed and then broken by sonication at 4 °C for 30 min. The cells were centrifuged at high speed for 15 min, and the supernatant was obtained and purified as previously described(Zhou et al., 2019). Briefly, the supernatant was loaded onto a Ni-nitrilotriacetic acid (NTA) column. The column was washed with washing buffer consisting of 0.5 M NaCl, 20 mM sodium phosphate, and 20 mM imidazole, pH 7.4. Then, the BmMETTL3(204-329) protein were eluted with a stepwise gradient of imidazole (up to 500 mM) in washing buffer. The eluate was dialyzed against 20 mM sodium phosphate buffer (pH 7.4). Subsequently, recombinant BmMETTL3(204-329) was stored at 4 °C or −80 °C until use.

### Preparation of antiserum and western blotting analysis

Purified proteins were used to generate rabbit polyclonal antibody according to previous study(Lee et al., 2016). Western blotting analysis was performed as previously described (Dai et al., 2019). In brief, Silkworm eggs and BmN cells were dissected and washed with cold phosphate-buffered saline (PBS). Furthermore, eggs and cells were mechanically homogenized in ice-cold RIPA lysate (RIPA: PMSF= 100: 1) (FDbio Science, China), incubated on ice for 1 h, and then centrifuged at 15,000 rpm for 15min at 4°C. The protein concentration in the supernatant (extracted protein) was estimated using the BCA method. The equal amount of protein (15μg) was separated by SDS-PAGE and transferred onto PVDF membranes (Millipore USA). After blocking with 5% nonfat milk for 2 h, the membranes were incubated overnight with polyclonal rabbit anti-BmMETTL3 immunoglobulin G. Subsequently, the membranes were incubated with a florescence-labeled secondary antibody (DingGuo ChangSheng Biotechnology Co., Ltd., China) for 2 h. The protein level of tubulin was used as a loading control. Western blotting signals were detected using an ECL Plus Kit (FDbio Science, China).

### Construction and transfection of plasmids for subcellular localization and overexpression

The ORFs encoding METTL3, METT14, FL2D, YTHDC and YTHDF3 were amplified and cloned into pIZ/V5-EGFP and pIZ/V5-His vector using specific primers list in Table S1. All plasmids used were transfected into BmN cells using Lipo8000 transfection reagent (Beyotime, China) following the manufacturer’s protocol. After incubation for 48 h, cells for subcellular localization were washed with PBS, stained with Hoechst 33258 and observed by Zeiss LSM780 laser scanning confocal microscope.

### Injection of siRNA into silkworm eggs

Eggs of *B. mori* (N4 stain) were collected within 5 h after oviposition. The newly laid eggs were washed with tap water and left for 2–3 min. The floated eggs were transferred to glass slides and aligned in the same direction under a dissection microscopy. After eggs were dried for about 1 h at room temperature, they were bonded using an adhesive. *METTL3*-siRNA and Control-siRNA oligonucleotides were synthesized by Tsingke Hangzhou. The concentration of siRNA was adjusted to 4pm/μl. 1-5nl siRNA of control and *METTL3* was injected into the embryos prepared using a micro-injector. The injected eggs were incubated in a wet chamber at 25°C until hatching.

### Transcriptome sequencing

Sequencing was performed on BGISEQ-500 sequencer (Shenzhen, China). The sequencing data was filtered with SOAPnuke (v1.5.2)(Li et al., 2008). The clean reads were mapped to the reference genome (version 1.0; GenBank Accession Number: 7091) using HISAT2 (v2.0.4)(Kim et al., 2015). Bowtie2 (v2.2.5) was applied to align the clean reads to the reference coding gene set(Langmead and Salzberg, 2012). The expression level of gene was calculated by RSEM (v1.2.12)(Li and Dewey, 2011). The fragments per kilobase of transcript permillion (FPKM) were used to quantify the expression level.

### Detection of m6A content

m6A RNA Methylation Assay Kit (Fluorometric; Epigentek) was used to measure the m6A level in total RNA by MultiSkan FC microplate reader (Thermo Fisher, USA) according to the manufacturer’s protocol. Two hundred nanogram of total RNA was used per assay well. m6A in total RNA is specifically recognized and bound by the capture and detection antibodies. After fluorounce enhancer (Epigentek) and fluorounce developer (Epigentek) was added and the signal was measured in MultiSkan FC microplate reader (Thermo Fisher, USA) at 530/590 nm.

### Statistics analysis

Using Student’s t-test and analysis of variance to analyze the experimental data. NS p?0.05, * p < 0.05, ** p < 0.01, *** p < 0.005, ****P<0.0001, according to the student’s t-test. All values of the study are represented as mean ±SD or SEM from more than three independent experiments. Data were analyzed using commercial statistical analysis software (GraphPad Prism v8.0; GraphPad Software, La Jolla, California).

## Results

### The m6A related genes are conserved among insects

In order to reveal whether m6A pathway is widespread and pervasive among insects, we investigated the distribution of m6A related genes in 207 insects. We used the amino acid sequences from each of the m6A related genes of *B. mori*, *D. melanogaster*, *Megalopta genalis, Agrilus planipennis* and *B. tabaci* as query object and obtained 209 homologs of METTL3 from 199 insects, 206 homologs of METTL14 from 201 insects, 189 homologs of YTHDC from 187 insects, 218 homologs of YTHDF3 from 202 insects and 197 homologs of FL2D from 191 insects. we constructed a phylogenetic tree of insects and annotated the number of m6A related genes into it (Fig. 1). The number of m6A related genes were not significantly different among insects, indicating that the m6A regulatory genes in insects don’t exhibit large-scale expansion and deletion. Taken together, we concluded that the number of m6A related genes are highly conserved in insects.

**Fig. 1.**
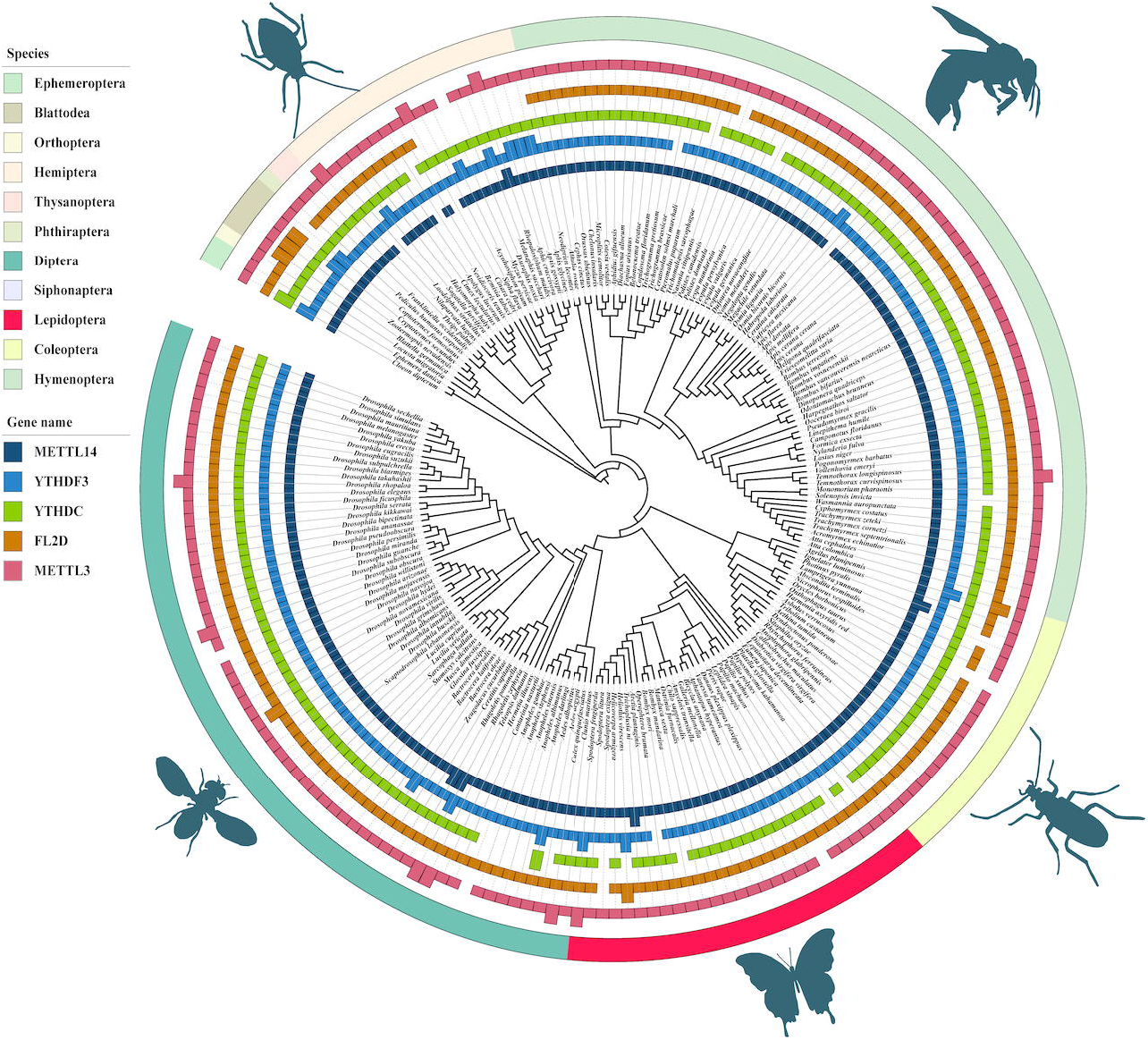
The number of each m6A related genes is detected by BLASTp and tBLASTn. This analysis indicated that the number of m6A genes in insects is conserved.

### Phylogenetic Analysis of METTL3

METTL3 catalyzes the formation of m6A which has important roles in regulating various biological processes(Xu et al., 2017). To elucidate the phylogenetic relationships of *METTL3* in silkworm and other insects, we reconstructed an ML phylogenetic tree with 209 full length *METTL3* amino acid sequences from 201 species. The result showed that *BmMETTL3* was much closer to *METTL3* of lepidoptera, and the *METTL3* of insects were, respectively, clustered, which was consistent with the evolutionary relationship of insect species (Fig.2). Subsequent multiple sequence alignments of the deduced amino acid sequences of *METTL3* from *Cloeon dipterum, Sitophilus oryzae, Dendroctonus ponderosae, Ephemera danica, Thrips palmi, Zootermopsis nevadensis, Frankliniella occidentalis, Blattella germanica, Cephus cinctus, Pogonomyrmex barbatus, Sipha flava, Acyrthosiphon pisum, B. mori, Galleria mellonella, D. melanogaster* and *Contarinia nasturtii* suggested that they have highly conserved MT-A70 domains (Fig S2). We also reconstructed the phylogenetic tree of other m6A related genes. The relationship of these m6A related genes as depicted by the phylogenic tree was generally in agreement with the traditional taxonomy of these species (Fig S3-Fig S6). These results illustrate that m6A related genes are evolutionarily conserved among insects.

**Fig. 2.**
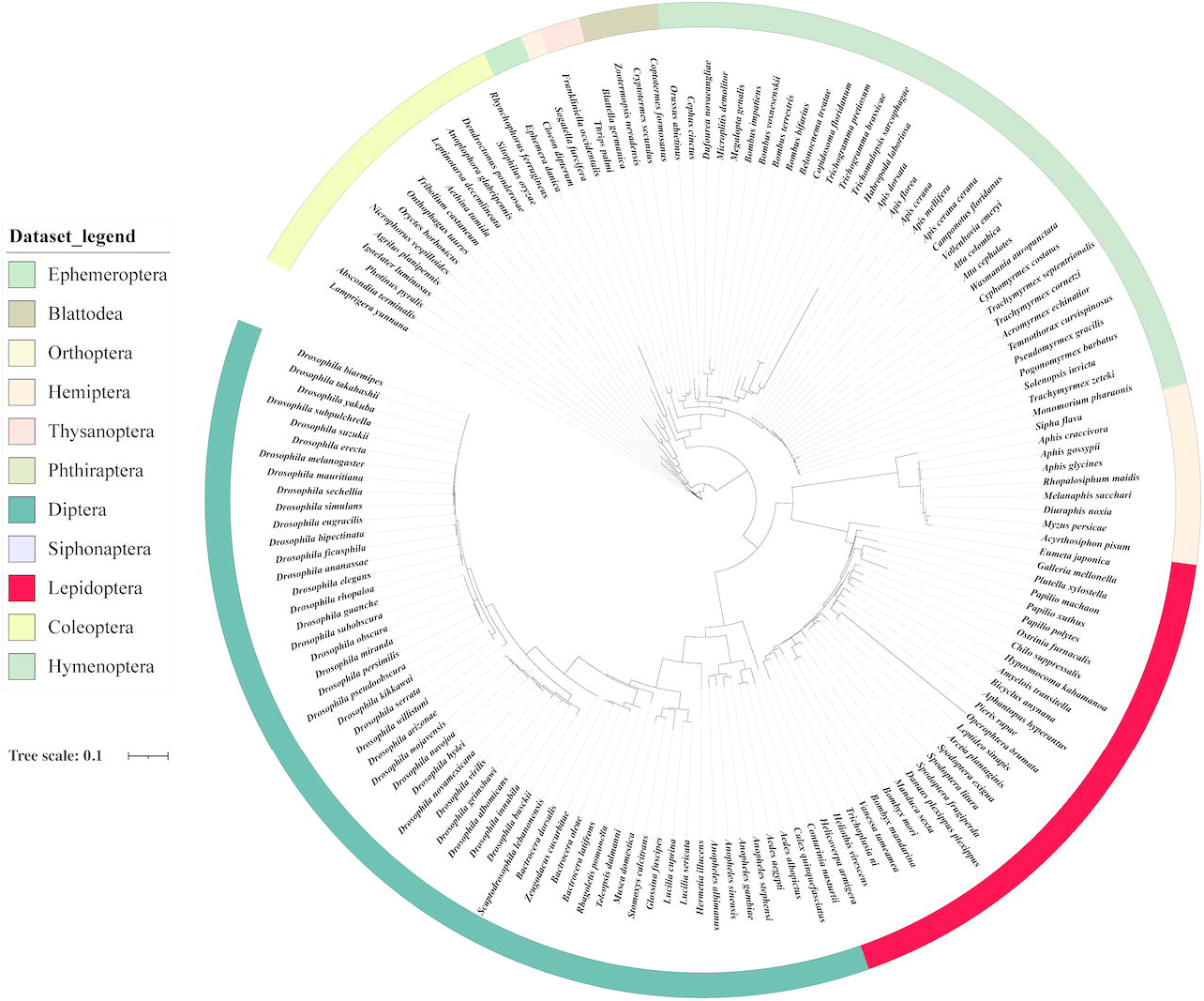
Phylogenetic analysis of *METTL3* in insects. The maximum-likelihood (ML) tree was constructed with MEGA 11.0.1 using the sequences amino acid sequences of 209 *METTL3* genes from 201 insect species.

### Subcellular localization of m6A related proteins in BmN cells

As subcellular localization is critical for functional analysis, we examined the subcellular localization of these m6A related proteins in BmN cells. we performed the transient expression assay. Green fluorescent protein (GFP)-tagged *METTL3, METTL14, FL2D, YTHDC* and *YTHDF3* was expressed in BmN cells. Confocal imaging showed that fluorescence signals of METTL3-GFP, METTL14-GFP and YTHDC-GFP fusion proteins precisely coincided with the nucleus in BmN cells (Fig.3, A1-C3). FL2D-GFP was distributed in both cytoplasm and nucleus with higher levels in the cytoplasm (Fig.3, D1-D3), while YTHDF3-GFP was found mainly in cytoplasm (Fig.3, E1-E3). These results illustrated that the subcellular location of m6A related proteins in BmN cells was consistent with mammalian cells (Uhlen et al., 2010; Lasman et al., 2020).

**Fig. 3.**
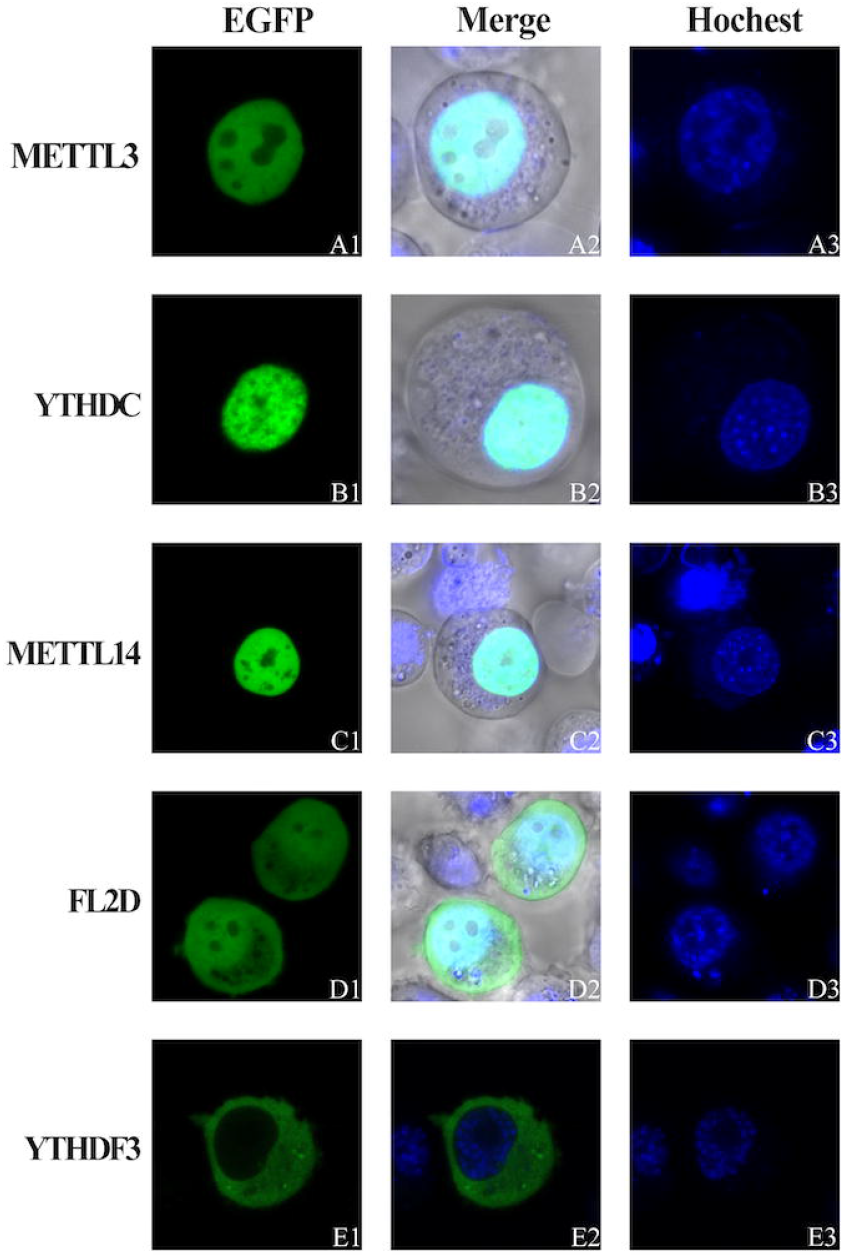
Expression of m6A related genes during *B.mori* embryonic development. Expression profiles of (*A)METTL14, (B)YTHDF3, (C)YTHDC, (D)FL2D* and (*E)METTL3* in eggs during the 8 day after laid. Data were normalized to *Bmrpl49* gene expression. (F)Identification of METTL3 protein by western blot.

### Expression patterns of m6A-related genes during B. mori embryonic development

To gain insights into how the dynamic m6A methyl marks are regulated during embryonic development of *B. mori*, we analyzed the expression profiles of m6A related genes by qPCR. *BmMETTL3* expression was observed at all stages examined, with particularly strong expression from Day6 to Day8 (Fig.4 A, F). *BmMETTL3* and *BmMETTL14*, as the main functional component of methyltransferase complex, have similar expression pattern (Fig.4 B). *BmFL2D*, which involved in the recruitment of *BmMETTL3* and *BmMETTL14*, has higher expression level than that of *BmMETTL3* and *BmMETTL14* (Fig.4 C). Reader proteins are both have high expression level in the first day. *BmYTHDF3* was highly expressed and the expression level fluctuates wildly from Day 2 to Day 8 compared to *BmYTHDC* (Fig.4 D, F). Taken together, m6A related genes were expressed widely throughout the embryo, and these genes showed different expression pattern during embryonic development of *B. mori*. The broad expression of m6A related genes during embryonic development suggests that it probably likely impact multiple developmental processes.

**Fig. 4.**
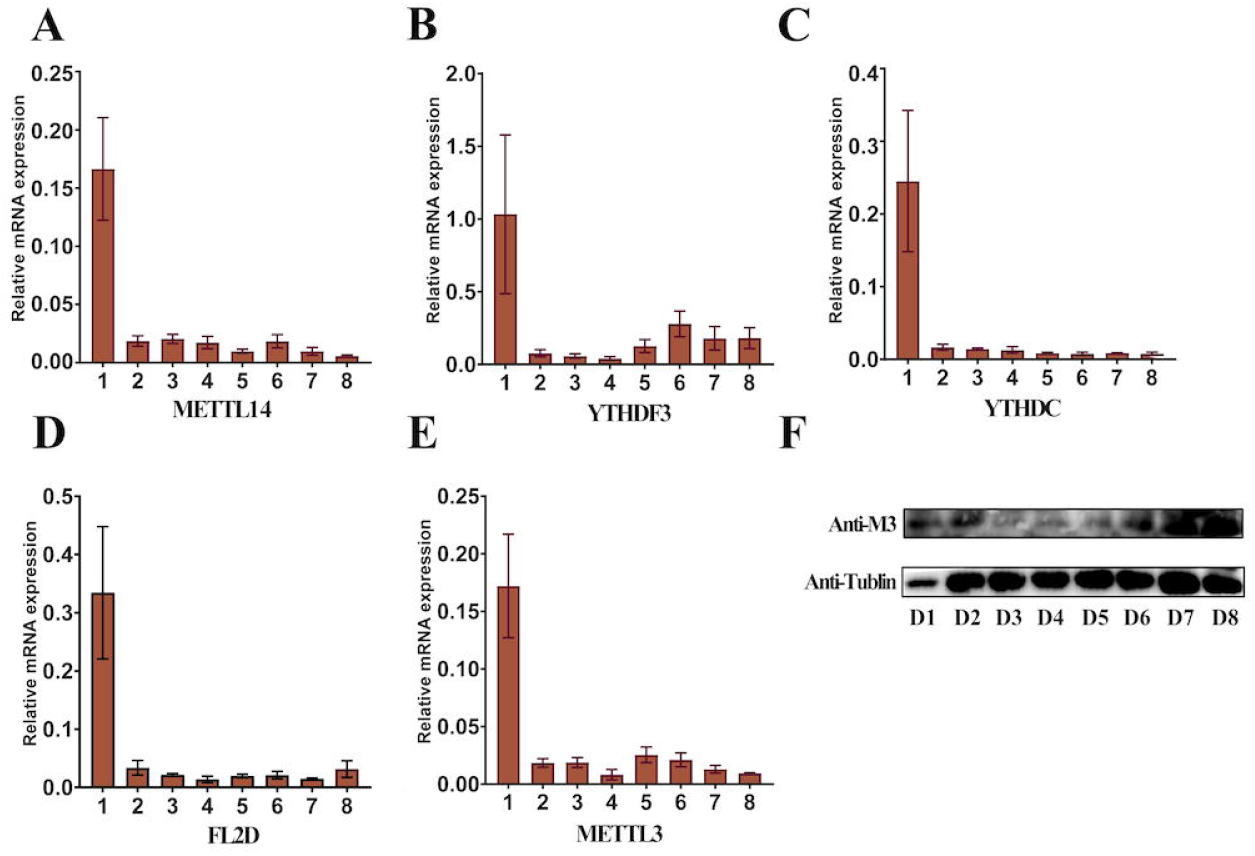
Subcellular localization of m6A reated proteins fused EGFP in BmN cells. The pIZ-EGFP-m6A related genes were transiently expressed in BmN cells. The cell nuclei were stained with Hoechst 33258 (blue). The fluorescent signal was imaged by confocal microscopy. EGFP green fluorescence, Hoechst 33258 blue fluorescence, and merged images are shown.

### Transcriptomic analysis of silkworm embryonic with suppressed METTL3 expression

*METTL3* is essential for embryonic development due to the critical role of m6A in timely RNA turnover (Batista et al., 2014). We further investigated the role of *METTL3* in *B. mori* embryonic development by microinjection of METTL3 siRNA. To determine the interference efficiency of METTL3 siRNA, we conducted Western blot analysis and qPCR after injection of METTL3 siRNA and control siRNA. METTL3 siRNA significantly reduced the expression of *METTL3* at 48 h (Fig.5 A, B). Moreover, the m6A level was significantly decreased in METTL3 siRNA group in comparison to control group (Fig.S7). The result is consistent with the function of METTL3, which as the methyltransferase core complex.

**Fig. 5.**
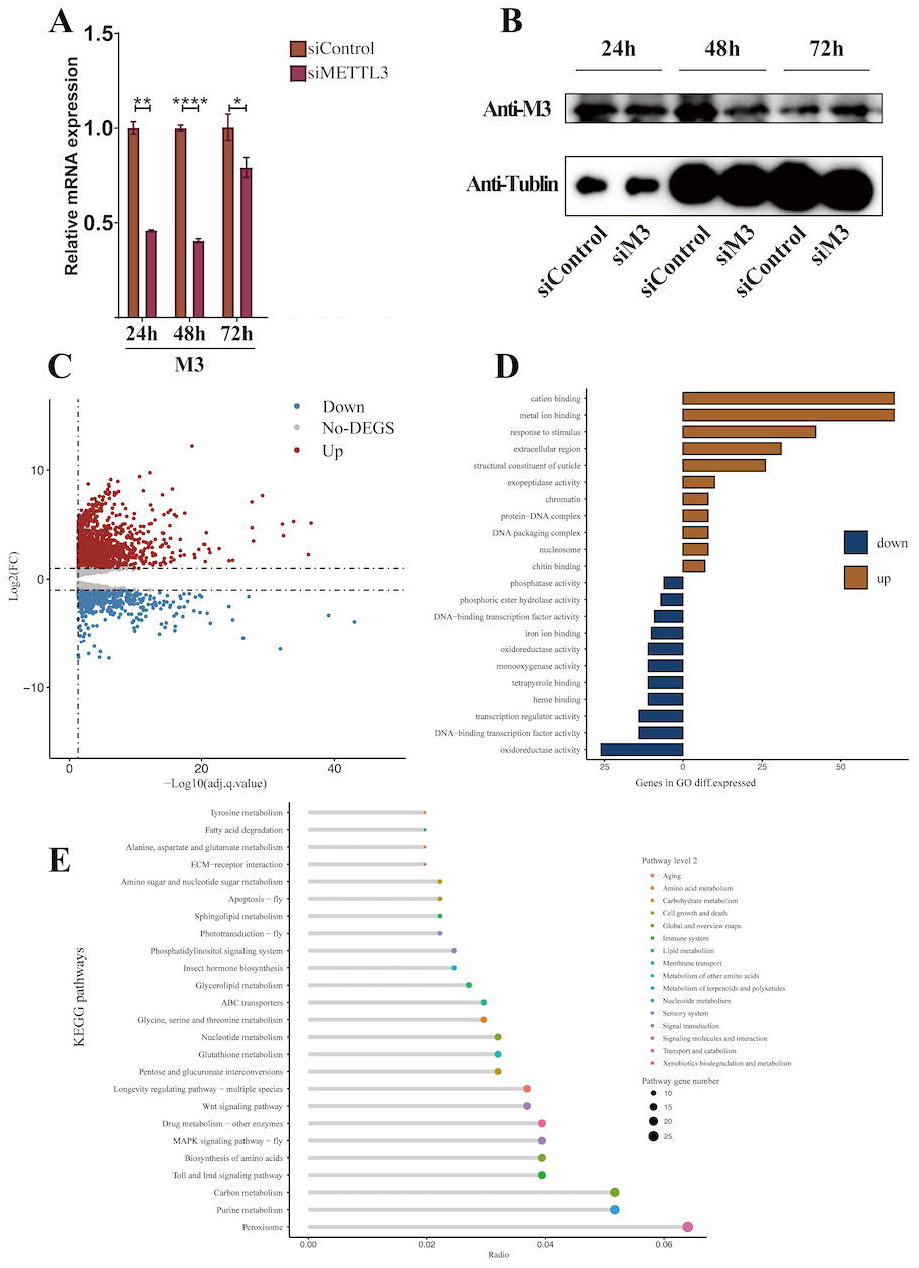
Transcriptomic analysis of silkworm embryonic with suppressed METTL3 expression. Expression levels of (A)*BmMETTL3* were quantified by quantitative RT-qPCR and normalized by the expression levels of *Bmrpl49*, (B) the *BmMETTL3* protein expression was measured by Western blotting. Relative expression levels with a negative control (negative siRNA) were represented. (C) The volcano plot of DGEs. Red spots indicate significantly up-regulated genes; blue spots indicate significantly down-regulated genes; grey spots are genes of no different expression. The significant difference is set at FDR <0.05 to identify the different expression genes between two groups. (D) KEGG pathway enrichment analysis of the DEGs showed the top 20 enriched KEGG terms. Different colors represent the level 2 of KEGG pathways. The size of point represents the numbers of enriched genes in pathways. (E) Enriched distributions of DEGs into GO categories according to GO enrichment analysis. Red bar are upregulated DEGs; blue bar are downregulated DEGs.

We further performed the transcriptomic analysis to identify the impact of METTL3 knockdown. Sequencing of RNA isolated from control siRNA and METTL3 siRNA injected silkworm eggs identified 2183 differentially expressed genes, of which 1528 were up-regulated in METTL3 siRNA treated eggs (Fig.5 C). GO enrichment analysis was performed for the up-regulated and down-regulated genes separately (Fig.5 D). In the downregulated group, enriched terms include DNA-binding transcription factor activity, RNA polymerase II-specific (GO: 0000981), DNA-binding transcription factor activity (GO:0003700), oxidoreductase activity (GO:0016491), and phosphatase activity (GO:0016791), etc. In the unregulated group enriched terms include cation binding (GO:0043169), Metal binding (GO:0046872), response to stimulus (GO:0050896), and extracellular region (GO:0005576), etc. GO functional categories of up-regulated genes enrichment include molecular function (MF), biological process (BP), and cellular component (CC). The enriched biological pathways by the DEGs were also analyzed with KEGG (Fig.5 E). The enriched KEGG pathways included peroxisome, purine metabolism, carbon metabolism, Toll and Imd signaling pathway, MAPK signaling pathway, Wnt signaling pathway, etc. These pathways include biochemical reactions, signaling pathways, and metabolic pathways that are important for silkworm embryonic development. These results indicate that *METTL3* silencing broadly affect biological processes including immune related, cellular homeostasis, organic substance metabolic process, biosynthetic process, energy metabolism and catabolism.

### BmMETTL3 knockdown affect Toll/Imd, Wnt signaling pathway activity

Toll/Imd and Wnt signaling pathway are essential for silkworm embryogenesis. Here, we found that *METTL3* silencing significantly affected several genes in Toll/Imd and Wnt signaling pathways. 11 genes were significantly up-regulated, and 8 genes were down-regulated in the Wnt signaling pathway (Fig.6A). The expression of *Wnt-4* and *PLCb1* were significantly upregulated 32-fold and 8-fold, respectively. The expression of Wnt-10b, Wnt inhibitory factor 1 was significantly downregulated for 2-fold after *METTL3* knock down. 9 genes were significantly up-regulated, and 9 genes were down-regulated in the Toll/Imd signaling pathway (Fig.6B). The expression of *PGRP* and peptidoglycan-recognition protein LB were significantly upregulated for 64-fold and 45-fold, respectively. The expression of cecropin-D-like peptide was significantly downregulated for 64-fold. These results suggested that m6A methylation significantly affected Toll/Imd and Wnt signaling pathway.

**Fig. 6.**
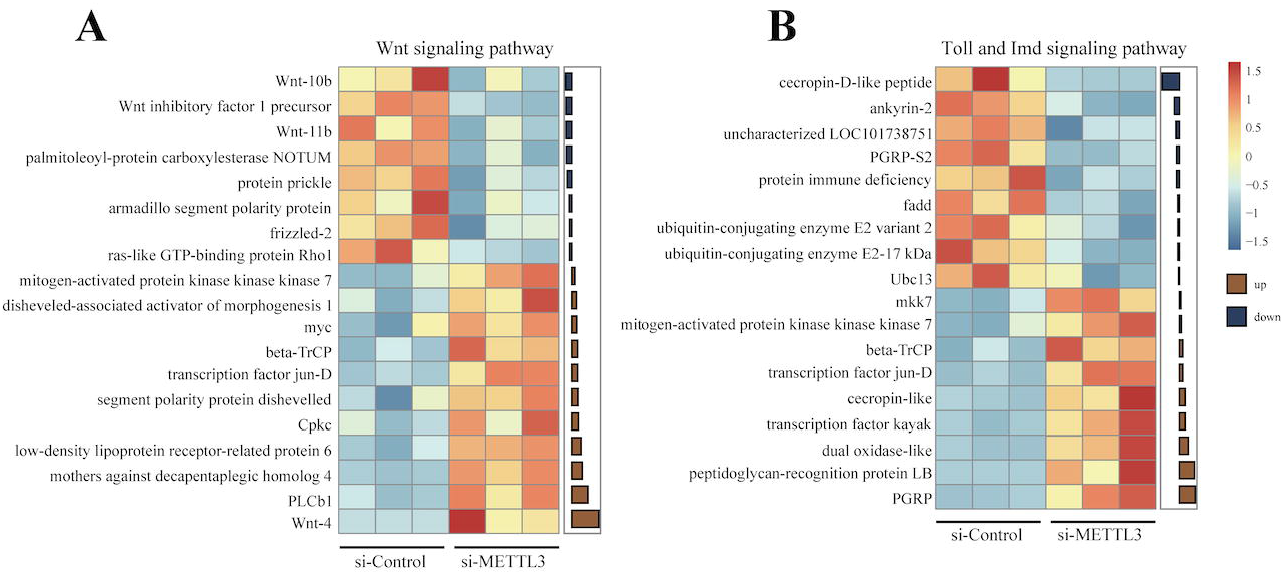
Analysis of differential genes involved in Wnt and Toll/Imd signaling pathways. Heat map of differentially expressed genes on the (A) Wnt signaling pathway and (B) Toll/Imd signaling pathway in *B. mori* embryonic with *METTL3* knockdown. Heat map colors correspond to the scale value of FPKM. Bar plots display the fold change (log2FC) of RNAi versus control.

### Overexpression of BmMETTL3 in BmN cells

To further verify the biological role of METTL3, BmMETTL3 was overexpressed in BmN cell lines. The qPCR and Western blot results showed that the expression of BmMETTL3 was significantly increased (Fig.7 A). We then performed genome-wide transcriptome profiling (RNA-seq) analysis in BmMETTL3-overexpressing BmN cells. A total of 320 DEGs were obtained between the oe-METTL3 group and control group with FDR<0.05, of which 163 (50.9%) were up-regulated and 157 (49.1%) were down-regulated (Fig.7 B). Clustering analysis showed that the expression patterns of DEGs were obviously different between the control group and METTL3-overexpression group.

**Fig. 7.**
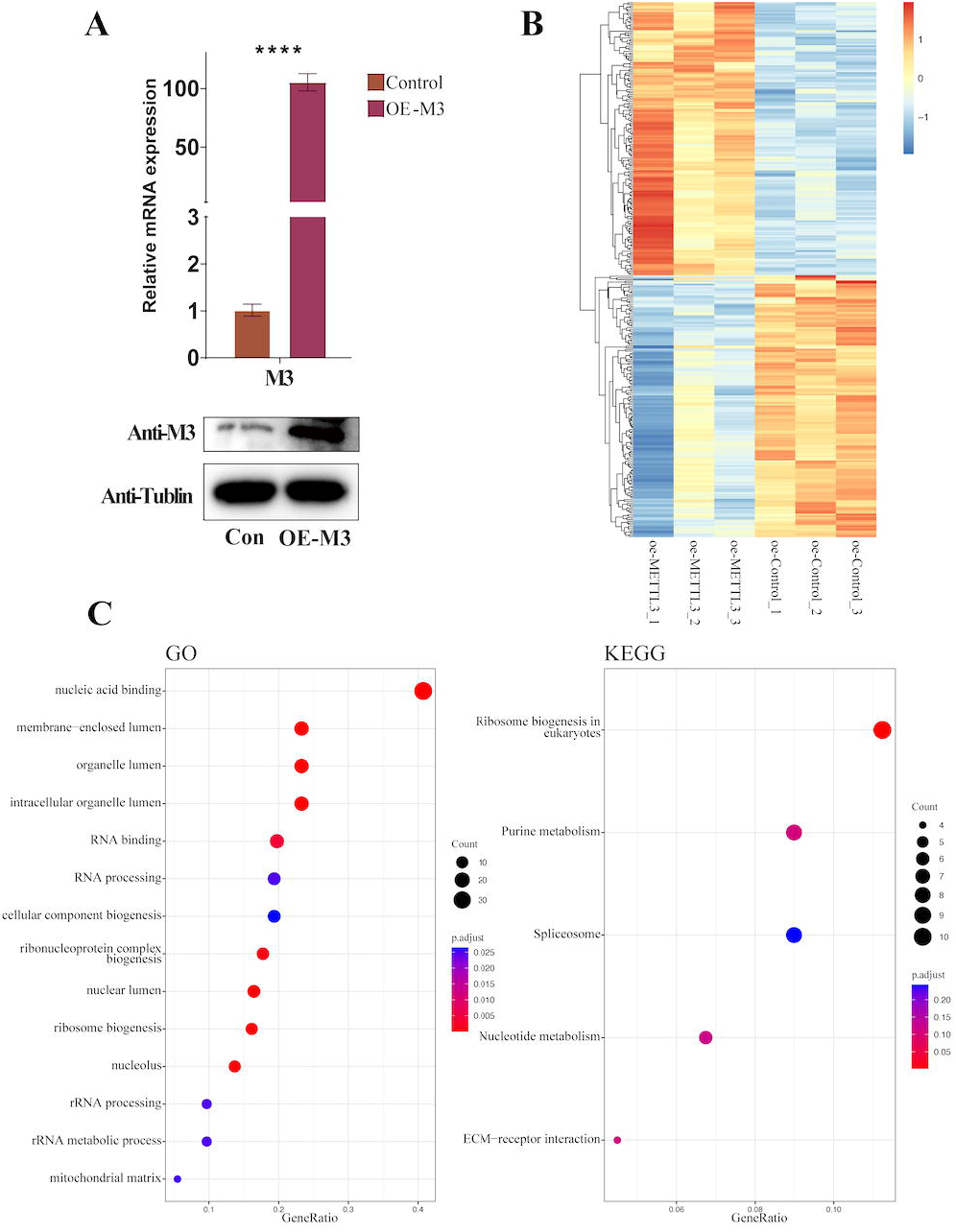
Transcriptomic analysis of BmN cells with BmMETTL3 overexpression. (A)Expression levels of BmMETTL3 were quantified by quantitative RT-PCR and Western blot after overexpression of BmMETTL3. (B) Heatmap analysis of differential genes. (C) GO and KEGG pathway enrichment analysis of the DEGs.

Based on GO annotation, GO terms nucleic acid binding (GO:0003676), RNA binding (GO:0003723), RNA processing (GO:0006396) were enriched in DEGs. The result is consistent with the role of METTL3 as a methyltransferase for m6A. we also found that GO terms nuclear lumen (GO:0031981), intracellular organelle lumen (GO:0070013), organelle lumen (GO:0043233) and membrane-enclosed lumen (GO:0031974) were also enriched, suggesting that BmMETTL3 overexpression affect organelle function (Fig.7 C).

The enrichment analysis of KEGG pathways show that DEGs were enriched in 5 pathways, including ribosome biogenesis in eukaryotes, purine metabolism, spliceosome, nucleotide metabolism and ECM-receptor interaction (Fig.7 C). Moreover, many DEGs related to proliferation, such as frizzled-2, tyrosine-protein kinase Dnt, G1/S-specific cyclin-D3 were also enriched in Wnt signaling pathway. Compared to the results that knockdown METTL3 in silk embryonic, overexpression of METTL3 also impact the nucleic acid binding, which have a significant influence of on nucleic acid metabolism. KEGG pathways analysis revealed that purine metabolism pathway was also affected by METTL3 overexpression. Together, the results show that METTL3 overexpression significantly influence nucleotide metabolism and organelle function in BmN cells.

## Discussion

m6A has been intensely studied in the dipteran *D. melanogaster* (Guo et al., 2018; Haussmann et al., 2016; Lence et al., 2016). The evolutionary relationships of m6A related genes and function of m6A in other insects, however, remain largely unknown. Here we carried out evolutionary analysis of m6A related genes in insects, and investigated the function of METTL3 in the embryonic development of lepidopteran silkworm. Our analysis suggested that the evolutionary relationship of m6A related genes is highly conserved in insects and METTL3 plays an important role in the regulation of Toll/Imd and Wnt signaling pathways and then influences the development of silkworm embryonic.

The evolutionary conservation of m6A modifications was detected within many species, including yeast (*Saccharomyces mikatae* and *Saccharomyces Cerevisiae*)(Schwartz et al., 2013); human (*Homo sapiens*), chimpanzee (*Pan troglodytes*), and rhesus (*Macaca mulatta*) (Ma et al., 2017). In this study we found that the number of m6A related genes are conserved in insects and the cellular localization of m6A related proteins are basically the same in mammals and *B. mori*. Our results show that the evolution of the m6A genes is mostly in accordance with the known phylogeny of the insects. The amino acid sequences of METTL3 and METTL14 alignment showed that the MT-A70 domains were highly conserved in insects. These results illustrate that these m6A methyltransferases are evolutionarily conserved among insect species.

The importance of m6A methylation has been clarified in multiple cellular processes of differentiation, self-renewal, and pluripotency during embryonic development (Sánchez-Vásquez et al., 2018). In *D. melanogaster*, m6A was remarkably enriched in early embryogenesis and the components of methyltransferases were also ubiquitously expressed in concert with these stages (Lence et al., 2016). Consistently, we showed a wide expression of m6A related genes throughout embryogenesis of *B. mori*. METTL3 is a key component of the m6A methyltransferase complex (Shi et al., 2019). In this study, our transcriptome analysis demonstrated that the knockdown of *BmMETTL3* induced a significant upregulation in DEGs that were functionally associated with cation binding, metal ion binding. By contrast, the extensive number in DEGs encoding enzymes for transcription regulator activity were largely downregulated.

The destabilization of m6A-containing mRNAs was first identified in studies that uncovered an increase in the half-life of mRNAs after m6A writer protein downregulation in both human and mouse cells. In this study, we found that the number of up-regulated genes was significantly more than the down-regulated genes (1528 vs. 655) for DEGs. The result might relevant to that m6A modification affect mRNA degradation.

In the study of *Anser cygnoides orientalis* embryos, DMGs (different m6A methylation genes) mainly enriched in Wnt signaling pathway, mTOR signaling pathway, and FoxO signaling pathway in KEGG pathway analysis(Xu et al., 2021). In our study, we also found that part of DEGs after *METTL3* knock down were enriched in Wnt signaling pathway. However, we found DEGs also enriched in MAPK, Toll/Imd and longevity regulating pathways, etc.

The Toll/Imd and Wnt signaling pathways are critical signal transduction pathway involved in the regulation of embryonic development. Wnt signaling has been shown to control diverse biological processes, including embryonic development, tissue regeneration, hematopoiesis, survival, cellular proliferation, and differentiation(Clevers, 2006; MacDonald et al., 2009; Prakash and Swaminathan, 2015). Recent studies reported that m6A affect a diverse array of physiologic activities in many cell types via regulation of Wnt signaling pathway. The reduction of RNA m6A methylation activated oncogenic Wnt/PI3K-Akt signaling, and promoted malignant phenotypes of GC cells(Zhang et al., 2019). Toll/Imd pathway is one of important signaling pathways regulating insect innate immunity. Our findings demonstrate that *METTL3* knockdown resulted in differential expression of genes in Wnt and Toll/Imd signaling pathway, and then affect the activity of the Wnt and Toll/Imd signaling pathways in *B. mori* embryonic.

Frizzled-2 is the Fzd2 gene product, Wnt5a regulates distinct signaling pathways by binding to Frizzled-2(Sato et al., 2010). G1/S-specific cyclin-D3 is essential regulator of G1/S transition in response to growth factor stimulation and a downstream target gene of Wnt/β-catenin pathway(Pang et al., 2017). They are both important component in Wnt signaling pathway. Both in vivo and in vitro experiments demonstrated that METTL3 regulates the expression of Frizzled-2 and G1/S-specific cyclin-D3.

Purine is precursor for the synthesis of primary and secondary products as well as the building blocks for nucleic acids, and purine metabolism is important component in the whole DNA synthesis process. In this study, we found that purine metabolism is an important target pathway for METTL3 in vivo and in vitro experiment.

Here, we performed a comprehensive analysis of the evolution of m6A related genes in insects. These genes have high sequence similarity, implying that insect m6A related genes were conserved throughout evolution. To the best of our knowledge, this is the first study to comprehensively ascertain the physiological functions of METTL3 in the embryonic development of insects. Our data illustrate that METTL3 influences a number of cellular pathways that are involved in development of embryonic including proliferation and metabolism. By focusing our analyses on Wnt and Toll/Imd signaling pathways, we identified METTL3 knockdown significantly affect both of the two pathways and then, influence embryonic development, tissue homeostasis and innate immunity. Our study has provided significant new insights into the physiologic function of METLL3 in insect embryos.

## Acknowledgements

This work was supported by the National Natural Science Foundation of China (Grant No. 31970460; 32170483), the Natural Science Foundation of Zhejiang Province (Grant No. LR22C040001), the Zhejiang Provincial Science and Technology Plans (2021C02072-6), and the Zhejiang Provincial Key Laboratory Construction Plans (2020E10025).

## Disclosure

The authors declare no conflict of interest with the contents of this article.

## Notes

### Competing Interest Statement

The authors have declared no competing interest.

