## Supplementary material for "Evolution of m6A related genes in insects and function of METTL3 in embryonic development of silkworm": Supplementary information.docx

**SUPPLEMENTAL INFORMATION**

**Table S1 List of primers used in this paper**

| **Primers** | **Sequence (5'--3')** | **Purpose** |
| --- | --- | --- |
| BmorMETTL3-F | CCCTATGGCACTATGTCGG | qRT-PCR of *BmMETTL3* |
| BmorMETTL3-R | CGGTTCTGATGATGCGTTG | qRT-PCR of *BmMETTL3* |
| BmorMETTL14-F | GTGTCTGGAAGCGGAGGTGA | qRT-PCR of *BmMETTL14* |
| BmorMETTL14-R | GGATTTGATGACTGCGTGCC | qRT-PCR of *BmMETTL14* |
| BmorYTHDF3-F | AACAGCGGCATTTGGACAAC | qRT-PCR of *BmYTHDF3* |
| BmorYTHDF3-R | GGCATACCAGGCCCTTTCTT | qRT-PCR of *BmYTHDF3* |
| BmorYTHDC-F | ACGGGATCGCTTTCCAATGT | qRT-PCR of *BmYTHDC* |
| BmorYTHDC-R | ATCGATCAGCAGTTCGCCAT | qRT-PCR of *BmYTHDC* |
| BmorFL2D-F | GTGACACGAACTCCATCCGA | qRT-PCR of *BmFL2D* |
| BmorFL2D-R | GCCTTTTCCAAGCCTCCTTC | qRT-PCR of *BmFL2D* |
| BmorRPL49-F | CAGGCGGTTCAAGGGTCAATAC | qRT-PCR of *BmRPL* |
| BmorRPL49-R | TGCTGGGCTCTTTCCACGA | qRT-PCR of *BmRPL* |
| pIZ-EGFP-BmMETTL3-F | ctcggtaccgagctcggatccATGTCGGACGCTTGGGAGGAAATA | Subcellular localization of BmMETTL3 |
| pIZ-EGFP-BmMETTL3-R | gcccttgctcaccattctagaAGTGAGTCCCGGGTCTGGCGGCGG | Subcellular localization of BmMETTL3 |
| pIZ-EGFP-BmMETTL14-F | ctcggtaccgagctcggatccATGAGTGAGAAATTAAGAGAACTA | Subcellular localization of BmMETTL14 |
| pIZ-EGFP-BmMETTL14-R | gcccttgctcaccattctagaGATAGCCCCTCTGCCCCTC | Subcellular localization of BmMETTL14 |
| pIZ-EGFP-BmFL2D-F | ctcggtaccgagctcggatccATGAGTGCAATGTCCGAGGAGA | Subcellular localization of BmFL2D |
| pIZ-EGFP-BmFL2D-R | gcccttgctcaccattctagaCTCCGCGTCGCCATTTGCGAC | Subcellular localization of BmFL2D |
| pIZ-EGFP-BmYTHDF3-F | ctcggtaccgagctcggatccATGTCAGCAGGCGTGTCAGAT | Subcellular localization of BmYTHDF3 |
| pIZ-EGFP-BmYTHDF3-R | gcccttgctcaccattctagaATTGCGGGGACGTCCTCG | Subcellular localization of BmYTHDF3 |
| pIZ-EGFP-BmYTHDC-F | ctcggtaccgagctcggatccATGGAGACCTCTAACAACACAGA | Subcellular localization of BmYTHDC |
| pIZ-EGFP-BmYTHDC-R | gcccttgctcaccattctagaGCGTCGATCACGATAAGACCTGCTT | Subcellular localization of BmYTHDC |
| pIZ-His-BmMETTL3-F | ctcggtaccgagctcggatccATGTCGGACGCTTGGGAGG | Over expression of BmMETTL3 |
| pIZ-His-BmMETTL3-R | gtgatgatgaccggtacgcgtAGTGAGTCCCGGGTCTGGC | Over expression of BmMETTL3 |
| siBmMETTL3-sense | CCTGCAAAAAGTTACATTT | RNA interference of BmMETTL3 |
| siBmMETTL3-antisense | AAATGTAACTTTTTGCAGG | RNA interference of BmMETTL3 |
| siControl-antisense | UUCUCCGAACGUGUCACGUTT | RNA interference of negative |
| siControl-antisense | ACGUGACACGUUCGGAGAATT | RNA interference of negative |

**Table S2 List of tree node label, species and accession number of YTHDF3 gene in insects**

| **Tree node label** | **Accession number** | **Species** |
| --- | --- | --- |
| Bombyx_mori | XP_004924605.1 | *Bombyx mori* |
| Manduca_sexta | XP_030027669.1 | *Manduca sexta* |
| Helicoverpa_armigera | XP_021182967.1 | *Helicoverpa armigera* |
| Arctia_plantaginis | CAB3220860.1 | *Arctia plantaginis* |
| Papilio_machaon | XP_014360510.1 | *Papilio machaon* |
| Trichoplusia_ni | XP_026726299.1 | *Trichoplusia ni* |
| Spodoptera_litura | XP_022818611.1 | *Spodoptera litura* |
| Spodoptera_frugiperda | XP_035429459.1 | *Spodoptera frugiperda* |
| Aphantopus_hyperantus | XP_034840899.1 | *Aphantopus hyperantus* |
| Vanessa_tameamea | XP_026484438.1 | *Vanessa tameamea* |
| Bicyclus_anynana | XP_023955064.1 | *Bicyclus anynana* |
| Amyelois_transitella | XP_013193183.1 | *Amyelois transitella* |
| Pieris_rapae | XP_022130163.1 | *Pieris rapae* |
| Galleria_mellonella | XP_026764097.1 | *Galleria mellonella* |
| Hyposmocoma_kahamanoa | XP_026323400.1 | *Hyposmocoma kahamanoa* |
| Chilo_suppressalis | RVE46905.1 | *Chilo suppressalis* |
| Papilio_xuthus | KPI93494.1 | *Papilio xuthus* |
| Leptidea_sinapis | VVC93036.1 | *Leptidea sinapis* |
| Danaus_plexippus_plexippus | XP_032530266.1 | *Danaus plexippus plexippus* |
| Ostrinia_furnacalis | XP_028164456.1 | *Ostrinia furnacalis* |
| Eumeta_japonica | GBP41588.1 | *Eumeta japonica* |
| Operophtera_brumata | KOB72287.1 | *Operophtera brumata* |
| Anoplophora_glabripennis | XP_018573803.1 | *Anoplophora glabripennis* |
| Leptinotarsa_decemlineata | XP_023014103.1 | *Leptinotarsa decemlineata* |
| Tribolium_castaneum | XP_015835734.1 | *Tribolium castaneum* |
| Photinus_pyralis | XP_031343886.1 | *Photinus pyralis* |
| Onthophagus_taurus | XP_022913685.1 | *Onthophagus taurus* |
| Nicrophorus_vespilloides | XP_017768674.1 | *Nicrophorus vespilloides* |
| Halyomorpha_halys | XP_014279832.1 | *Halyomorpha halys* |
| Abscondita_terminalis | KAF5298474.1 | *Abscondita terminalis* |
| Thrips_palmi | XP_034232176.1 | *Thrips palmi* |
| Agrilus_planipennis | XP_018319502.1 | *Agrilus planipennis* |
| Diabrotica_virgifera_virgifera | XP_028134070.1 | *Diabrotica virgifera virgifera* |
| Frankliniella_occidentalis | KAE8741864.1 | *Frankliniella occidentalis* |
| Cimex_lectularius | XP_014244077.1 | *Cimex lectularius* |
| Neodiprion_lecontei | XP_015516544.1 | *Neodiprion lecontei* |
| Rhynchophorus_ferrugineus | KAF7281367.1 | *Rhynchophorus ferrugineus* |
| Aethina_tumida | XP_019876443.1 | *Aethina tumida* |
| Apolygus_lucorum | KAF6204476.1 | *Apolygus lucorum* |
| Athalia_rosae | XP_012264071.1 | *Athalia rosae* |
| Cryptotermes_secundus | XP_023713789.1 | *Cryptotermes secundus* |
| Sitophilus_oryzae | XP_030749638.1 | *Sitophilus oryzae* |
| Dendroctonus_ponderosae | XP_019761491.1 | *Dendroctonus ponderosae* |
| Zootermopsis_nevadensis | XP_021924743.1 | *Zootermopsis nevadensis* |
| Ctenocephalides_felis | XP_026466905.1 | *Ctenocephalides felis* |
| Polistes_dominula | XP_015172663.1 | *Polistes dominula* |
| Cephus_cinctus | XP_024937244.1 | *Cephus cinctus* |
| Odontomachus_brunneus | XP_032664254.1 | *Odontomachus brunneus* |
| Vespa_mandarinia | XP_035734692.1 | *Vespa mandarinia* |
| Lamprigera_yunnana | KAF5295403.1 | *Lamprigera yunnana* |
| Vespula_germanica | KAF7406542.1 | *Vespula germanica* |
| Camponotus_floridanus | XP_019885326.2 | *Camponotus floridanus* |
| Vespula_pensylvanica | KAF7430105.1 | *Vespula pensylvanica* |
| Vollenhovia_emeryi | XP_011864579.1 | *Vollenhovia emeryi* |
| Cyphomyrmex_costatus | XP_018399044.1 | *Cyphomyrmex costatus* |
| Polistes_canadensis | XP_014615820.1 | *Polistes canadensis* |
| Blattella_germanica | PSN55172.1 | *Blattella germanica* |
| Harpegnathos_saltator | XP_025155561.1 | *Harpegnathos saltator* |
| Nylanderia_fulva | XP_029161274.1 | *Nylanderia fulva* |
| Ooceraea_biroi | XP_011349252.1 | *Ooceraea biroi* |
| Dinoponera_quadriceps | XP_014473440.1 | *Dinoponera quadriceps* |
| Formica_exsecta | XP_029679182.1 | *Formica exsecta* |
| Coptotermes_formosanus | GFG36390.1 | *Coptotermes formosanus* |
| Locusta_migratoria | JAMg_model_4175.1 | *Locusta migratoria* |
| Atta_cephalotes | XP_012063853.1 | *Atta cephalotes* |
| Trachymyrmex_septentrionalis | XP_018340831.1 | *Trachymyrmex septentrionalis* |
| Acromyrmex_echinatior | XP_011061675.1 | *Acromyrmex echinatior* |
| Apis_cerana_cerana | PBC25927.1 | *Apis cerana cerana* |
| Belonocnema_treatae | XP_033207520.1 | *Belonocnema treatae* |
| Wasmannia_auropunctata | XP_011708052.1 | *Wasmannia auropunctata* |
| Atta_colombica | XP_018057153.1 | *Atta colombica* |
| Cloeon_dipterum | CAB3386546.1 | *Cloeon dipterum* |
| Linepithema_humile | XP_012231705.1 | *Linepithema humile* |
| Trachymyrmex_zeteki | XP_018314222.1 | *Trachymyrmex zeteki* |
| Trachymyrmex_cornetzi | XP_018370694.1 | *Trachymyrmex cornetzi* |
| Apis_cerana | XP_016904666.1 | *Apis cerana* |
| Apis_florea | XP_003693800.1 | *Apis florea* |
| Apis_mellifera | XP_624228.1 | *Apis mellifera* |
| Apis_dorsata | XP_006612493.1 | *Apis dorsata* |
| Chelonus_insularis | XP_034937363.1 | *Chelonus insularis* |
| Pogonomyrmex_barbatus | XP_011634790.1 | *Pogonomyrmex barbatus* |
| Osmia_bicornis_bicornis | XP_029040431.1 | *Osmia bicornis bicornis* |
| Osmia_lignaria | XP_034183853.1 | *Osmia lignaria* |
| Trichogramma_pretiosum | XP_014225752.1 | *Trichogramma pretiosum* |
| Temnothorax_curvispinosus | XP_024880082.1 | *Temnothorax curvispinosus* |
| Ignelater_luminosus | KAF2900988.1 | *Ignelater luminosus* |
| Habropoda_laboriosa | XP_017790062.1 | *Habropoda laboriosa* |
| Eufriesea_mexicana | XP_017757754.1 | *Eufriesea mexicana* |
| Nasonia_vitripennis | XP_008208758.1 | *Nasonia vitripennis* |
| Diachasma_alloeum | XP_015126342.1 | *Diachasma alloeum* |
| Bombus_bifarius | XP_033307676.1 | *Bombus bifarius* |
| Bombus_terrestris | XP_003398397.1 | *Bombus terrestris* |
| Dufourea_novaeangliae | XP_015438457.1 | *Dufourea novaeangliae* |
| Bombus_impatiens | XP_003488431.1 | *Bombus impatiens* |
| Bombus_vancouverensis_nearcticus | XP_033193246.1 | *Bombus vancouverensis nearcticus* |
| Bombus_vosnesenskii | XP_033350063.1 | *Bombus vosnesenskii* |
| Orussus_abietinus | XP_012284706.1 | *Orussus abietinus* |
| Copidosoma_floridanum | XP_014208176.1 | *Copidosoma floridanum* |
| Melipona_quadrifasciata | KOX78134.1 | *Melipona quadrifasciata* |
| Trichogramma_brassicae | CAB0032416.1 | *Trichogramma brassicae* |
| Monomorium_pharaonis | XP_036140817.1 | *Monomorium pharaonis* |
| Nomia_melanderi | XP_031830524.1 | *Nomia melanderi* |
| Callosobruchus_maculatus | VEN45739.1 | *Callosobruchus maculatus* |
| Megalopta_genalis | XP_033339578.1 | *Megalopta genalis* |
| Ceratina_calcarata | XP_017882313.1 | *Ceratina calcarata* |
| Pediculus_humanus_corporis | XP_002424976.1 | *Pediculus humanus corporis* |
| Fopius_arisanus | XP_011298104.1 | *Fopius arisanus* |
| Microplitis_demolitor | XP_008544641.1 | *Microplitis demolitor* |
| Pseudomyrmex_gracilis | XP_020295924.1 | *Pseudomyrmex gracilis* |
| Megachile_rotundata | XP_012148942.1 | *Megachile rotundata* |
| Ephemera_danica | KAF4526438.1 | *Ephemera danica* |
| Bemisia_tabaci | XP_018895647.1 | *Bemisia tabaci* |
| Temnothorax_longispinosus | TGZ37519.1 | *Temnothorax longispinosus* |
| Aphidius_gifuensis | KAF7996508.1 | *Aphidius gifuensis* |
| Frieseomelitta_varia | KAF3426044.1 | *Frieseomelitta varia* |
| Ceratosolen_solmsi_marchali | XP_011505010.1 | *Ceratosolen solmsi marchali* |
| Oryctes_borbonicus | KRT81681.1 | *Oryctes borbonicus* |
| Lasius_niger | KMQ98217.1 | *Lasius niger* |
| Clunio_marinus | CRK97438.1 | *Clunio marinus* |
| Rhopalosiphum_maidis | XP_026815524.1 | *Rhopalosiphum maidis* |
| Aphis_craccivora | KAF0749232.1 | *Aphis craccivora* |
| Aphis_glycines | KAE9527739.1 | *Aphis glycines* |
| Acyrthosiphon_pisum | XP_008180438.1 | *Acyrthosiphon pisum* |
| Aphis_gossypii | XP_027851368.1 | *Aphis gossypii* |
| Myzus_persicae | XP_022161443.1 | *Myzus persicae* |
| Melanaphis_sacchari | XP_025203937.1 | *Melanaphis sacchari* |
| Hermetia_illucens | XP_037922800.1 | *Hermetia illucens* |
| Sipha_flava | XP_025420367.1 | *Sipha flava* |
| Diuraphis_noxia | XP_015365166.1 | *Diuraphis noxia* |
| Drosophila_subpulchrella | XP_037709476.1 | *Drosophila subpulchrella* |
| Culex_quinquefasciatus | XP_001861474.1 | *Culex quinquefasciatus* |
| Cinara_cedri | VVC32945.1 | *Cinara cedri* |
| Drosophila_hydei | XP_023169635.2 | *Drosophila hydei* |
| Stomoxys_calcitrans | XP_013113692.1 | *Stomoxys calcitrans* |
| Contarinia_nasturtii | XP_031636297.1 | *Contarinia nasturtii* |
| Aedes_albopictus | XP_029729170.1 | *Aedes albopictus* |

**Table S3 List of tree node label, species and accession number of METTL3 gene in insects**

| **Tree node label** | **Accession number** | **Species** |
| --- | --- | --- |
| Bombyx_mori | XP_037874102.1 | *Bombyx mori* |
| Bombyx_mandarina | XP_028032854.1 | *Bombyx mandarina* |
| Spodoptera_litura | XP_022818540.1 | *Spodoptera litura* |
| Spodoptera_frugiperda | XP_035451896.1 | *Spodoptera frugiperda* |
| Manduca_sexta | XP_037296280.1 | *Manduca sexta* |
| Pieris_rapae | XP_022119130.1 | *Pieris rapae* |
| Hyposmocoma_kahamanoa | XP_026332810.1 | *Hyposmocoma kahamanoa* |
| Trichoplusia_ni | XP_026728190.1, XP_026746198.1 | *Trichoplusia ni* |
| Aphantopus_hyperantus | XP_034826507.1 | *Aphantopus hyperantus* |
| Chilo_suppressalis | RVE45703.1 | *Chilo suppressalis* |
| Bicyclus_anynana | XP_023947224.1 | *Bicyclus anynana* |
| Amyelois_transitella | XP_013191837.1 | *Amyelois transitella* |
| Vanessa_tameamea | XP_026499361.1 | *Vanessa tameamea* |
| Ostrinia_furnacalis | XP_028160989.1 | *Ostrinia furnacalis* |
| Heliothis_virescens | PCG81004.1 | *Heliothis virescens* |
| Arctia_plantaginis | CAB3229720.1 | *Arctia plantaginis* |
| Papilio_xuthus | KPJ04167.1 | *Papilio xuthus* |
| Galleria_mellonella | XP_026750524.1 | *Galleria mellonella* |
| Plutella_xylostella | XP_037965437.1 | *Plutella xylostella* |
| Spodoptera_exigua | KAF9423360.1 | *Spodoptera exigua* |
| Eumeta_japonica | GBP44075.1 | *Eumeta japonica* |
| Helicoverpa_armigera | XP_021198197.1 | *Helicoverpa armigera* |
| Danaus_plexippus_plexippus | XP_032524212.1 | *Danaus plexippus plexippus* |
| Ctenocephalides_felis | XP_026482233.1 | *Ctenocephalides felis* |
| Cryptotermes_secundus | XP_033610421.1 | *Cryptotermes secundus* |
| Blattella_germanica | PSN39413.1 | *Blattella germanica* |
| Papilio_polytes | XP_013145443.1 | *Papilio polytes* |
| Papilio_machaon | XP_014368296.1 | *Papilio machaon* |
| Onthophagus_taurus | XP_022907975.1 | *Onthophagus taurus* |
| Photinus_pyralis | XP_031328559.1, XP_031347075.1 | *Photinus pyralis* |
| Thrips_palmi | XP_034250139.1 | *Thrips palmi* |
| Zootermopsis_nevadensis | XP_021918306.1 | *Zootermopsis nevadensis* |
| Frankliniella_occidentalis | KAE8752518.1 | *Frankliniella occidentalis* |
| Ignelater_luminosus | KAF2885234.1 | *Ignelater luminosus* |
| Bemisia_tabaci | XP_018914444.1 | *Bemisia tabaci* |
| Abscondita_terminalis | KAF5299557.1 | *Abscondita terminalis* |
| Nicrophorus_vespilloides | XP_017773597.1 | *Nicrophorus vespilloides* |
| Coptotermes_formosanus | GFG29009.1 | *Coptotermes formosanus* |
| Pediculus_humanus_corporis | XP_002429182.1 | *Pediculus humanus corporis* |
| Agrilus_planipennis | XP_018327837.1 | *Agrilus planipennis* |
| Ephemera_danica | KAF4522221.1 | *Ephemera danica* |
| Asbolus_verrucosus | RZC39984.1 | *Asbolus verrucosus* |
| Cloeon_dipterum | CAB3361192.1 | *Cloeon dipterum* |
| Tribolium_castaneum | XP_967914.2 | *Tribolium castaneum* |
| Leptinotarsa_decemlineata | XP_023013922.1 | *Leptinotarsa decemlineata* |
| Halyomorpha_halys | XP_014271709.1 | *Halyomorpha halys* |
| Dendroctonus_ponderosae | XP_019772519.1 | *Dendroctonus ponderosae* |
| Hermetia_illucens | XP_037903744.1 | *Hermetia illucens* |
| Anoplophora_glabripennis | XP_018561753.1 | *Anoplophora glabripennis* |
| Callosobruchus_maculatus | VEN53699.1 | *Callosobruchus maculatus* |
| Rhynchophorus_ferrugineus | KAF7274373.1 | *Rhynchophorus ferrugineus* |
| Aethina_tumida | XP_019866496.1 | *Aethina tumida* |
| Sitophilus_oryzae | XP_030756474.1 | *Sitophilus oryzae* |
| Cimex_lectularius | XP_014262101.1 | *Cimex lectularius* |
| Aedes_albopictus | XP_019553766.2 | *Aedes albopictus* |
| Nasonia_vitripennis | XP_001606001.2 | *Nasonia vitripennis* |
| Anopheles_stephensi | XP_035907532.1 | *Anopheles stephensi* |
| Ceratosolen_solmsi_marchali | XP_011497733.1 | *Ceratosolen solmsi marchali* |
| Trichomalopsis_sarcophagae | OXU22029.1 | *Trichomalopsis sarcophagae* |
| Aedes_aegypti | XP_001662288.2 | *Aedes aegypti* |
| Camponotus_floridanus | XP_011253998.1 | *Camponotus floridanus* |
| Copidosoma_floridanum | XP_014203696.1 | *Copidosoma floridanum* |
| Anopheles_sinensis | KFB36575.1 | *Anopheles sinensis* |
| Wasmannia_auropunctata | XP_011694861.1 | *Wasmannia auropunctata* |
| Culex_quinquefasciatus | XP_001863984.1 | *Culex quinquefasciatus* |
| Monomorium_pharaonis | XP_012541939.1 | *Monomorium pharaonis* |
| Ceratitis_capitata | CAD7013394.1 | *Ceratitis capitata* |
| Polistes_dominula | XP_015190795.1 | *Polistes dominula* |
| Nylanderia_fulva | XP_029156687.1 | *Nylanderia fulva* |
| Ooceraea_biroi | XP_011339862.1 | *Ooceraea biroi* |
| Temnothorax_longispinosus | TGZ57472.1 | *Temnothorax longispinosus* |
| Trichogramma_brassicae | CAB0031542.1 | *Trichogramma brassicae* |
| Polistes_canadensis | XP_014614381.1 | *Polistes canadensis* |
| Trichogramma_pretiosum | XP_014237237.1 | *Trichogramma pretiosum* |
| Temnothorax_curvispinosus | XP_024884420.1 | *Temnothorax curvispinosus* |
| Vollenhovia_emeryi | XP_011874882.1 | *Vollenhovia emeryi* |
| Cyphomyrmex_costatus | XP_018407296.1 | *Cyphomyrmex costatus* |
| Pseudomyrmex_gracilis | XP_020285215.1 | *Pseudomyrmex gracilis* |
| Osmia_lignaria | XP_034172407.1 | *Osmia lignaria* |
| Chelonus_insularis | XP_034949048.1 | *Chelonus insularis* |
| Rhopalosiphum_maidis | XP_026822020.1, XP_026817490.1 | *Rhopalosiphum maidis* |
| Vespula_germanica | KAF7401701.1 | *Vespula germanica* |
| Vespula_pensylvanica | KAF7425673.1 | *Vespula pensylvanica* |
| Vespula_vulgaris | KAF7398879.1 | *Vespula vulgaris* |
| Bombus_impatiens | XP_012247962.2 | *Bombus impatiens* |
| Bombus_vosnesenskii | XP_033357057.1 | *Bombus vosnesenskii* |
| Acromyrmex_echinatior | XP_011057769.1 | *Acromyrmex echinatior* |
| Bombus_bifarius | XP_033307005.1 | *Bombus bifarius* |
| Bombus_vancouverensis_nearcticus | XP_033188546.1 | *Bombus vancouverensis nearcticus* |
| Atta_cephalotes | XP_012060234.1 | *Atta cephalotes* |
| Atta_colombica | XP_018060388.1 | *Atta colombica* |
| Trachymyrmex_septentrionalis | XP_018354296.1 | *Trachymyrmex septentrionalis* |
| Apis_dorsata | XP_006622662.1 | *Apis dorsata* |
| Osmia_bicornis_bicornis | XP_029033977.1 | *Osmia bicornis bicornis* |
| Bombus_terrestris | XP_012165914.2 | *Bombus terrestris* |
| Trachymyrmex_zeteki | XP_018304314.1 | *Trachymyrmex zeteki* |
| Odontomachus_brunneus | XP_032682345.1 | *Odontomachus brunneus* |
| Vespa_mandarinia | XP_035726029.1 | *Vespa mandarinia* |
| Bactrocera_oleae | XP_014088588.1 | *Bactrocera oleae* |
| Solenopsis_invicta | XP_011156250.1 | *Solenopsis invicta* |
| Eufriesea_mexicana | XP_017766012.1 | *Eufriesea mexicana* |
| Drosophila_yakuba | XP_002099307.1 | *Drosophila yakuba* |
| Megachile_rotundata | XP_012151596.1 | *Megachile rotundata* |
| Pogonomyrmex_barbatus | XP_011638177.1 | *Pogonomyrmex barbatus* |
| Nomia_melanderi | XP_031844697.1 | *Nomia melanderi* |
| Drosophila_erecta | XP_001982059.1 | *Drosophila erecta* |
| Drosophila_mauritiana | XP_033163214.1 | *Drosophila mauritiana* |
| Trachymyrmex_cornetzi | XP_018358216.1 | *Trachymyrmex cornetzi* |
| Drosophila_ficusphila | XP_017039820.1 | *Drosophila ficusphila* |
| Formica_exsecta | XP_029674888.1 | *Formica exsecta* |
| Aphis_gossypii | XP_027842849.1 | *Aphis gossypii* |
| Anopheles_gambiae | XP_312015.5 | *Anopheles gambiae* |
| Bactrocera_dorsalis | XP_011201082.1 | *Bactrocera dorsalis* |
| Megalopta_genalis | XP_033341118.1 | *Megalopta genalis* |
| Aphis_craccivora | KAF0770102.1 | *Aphis craccivora* |
| Harpegnathos_saltator | XP_011151071.1 | *Harpegnathos saltator* |
| Cephus_cinctus | XP_015594376.1 | *Cephus cinctus* |
| Acyrthosiphon_pisum | XP_001945512.1 | *Acyrthosiphon pisum* |
| Rhagoletis_pomonella | XP_036342182.1, XP_036343892.1 | *Rhagoletis pomonella* |
| Rhagoletis_zephyria | XP_017472883.1, XP_017490767.1 | *Rhagoletis zephyria* |
| Dufourea_novaeangliae | XP_015432967.1 | *Dufourea novaeangliae* |
| Drosophila_simulans | XP_002104643.2 | *Drosophila simulans* |
| Melanaphis_sacchari | XP_025203532.1 | *Melanaphis sacchari* |
| Drosophila_eugracilis | XP_017084184.1 | *Drosophila eugracilis* |
| Drosophila_kikkawai | XP_017036365.1 | *Drosophila kikkawai* |
| Oryctes_borbonicus | KRT80863.1 | *Oryctes borbonicus* |
| Drosophila_serrata | XP_020817780.1 | *Drosophila serrata* |
| Bactrocera_latifrons | XP_018799115.1 | *Bactrocera latifrons* |
| Orussus_abietinus | XP_012278176.1 | *Orussus abietinus* |
| Melipona_quadrifasciata | KOX70831.1 | *Melipona quadrifasciata* |
| Teleopsis_dalmanni | XP_037945373.1 | *Teleopsis dalmanni* |
| Apis_mellifera | XP_016767532.1 | *Apis mellifera* |
| Apis_cerana_cerana | PBC29210.1 | *Apis cerana cerana* |
| Apis_cerana | XP_016922633.1 | *Apis cerana* |
| Frieseomelitta_varia | KAF3427832.1 | *Frieseomelitta varia* |
| Sarcophaga_bullata | TMW42616.1 | *Sarcophaga bullata* |
| Drosophila_melanogaster | NP_651204.1 | *Drosophila melanogaster* |
| Drosophila_bipectinata | XP_017103172.1 | *Drosophila bipectinata* |
| Diuraphis_noxia | XP_015376446.1 | *Diuraphis noxia* |
| Diachasma_alloeum | XP_015123857.1 | *Diachasma alloeum* |
| Drosophila_biarmipes | XP_016952882.1 | *Drosophila biarmipes* |
| Drosophila_subpulchrella | XP_037717890.1 | *Drosophila subpulchrella* |
| Drosophila_sechellia | XP_032577660.1 | *Drosophila sechellia* |
| Microplitis_demolitor | XP_008560809.1 | *Microplitis demolitor* |
| Linepithema_humile | XP_012227311.1 | *Linepithema humile* |
| Zeugodacus_cucurbitae | XP_011178814.1 | *Zeugodacus cucurbitae* |
| Belonocnema_treatae | XP_033218805.1 | *Belonocnema treatae* |
| Drosophila_ananassae | XP_001953842.1 | *Drosophila ananassae* |
| Sipha_flava | XP_025411382.1 | *Sipha flava* |
| Apis_florea | XP_003697253.2 | *Apis florea* |
| Scaptodrosophila_lebanonensis | XP_030369246.1 | *Scaptodrosophila lebanonensis* |
| Athalia_rosae | XP_012259780.1 | *Athalia rosae* |
| Dinoponera_quadriceps | XP_014480930.1 | *Dinoponera quadriceps* |
| Lucilia_sericata | XP_037820919.1 | *Lucilia sericata* |
| Drosophila_persimilis | XP_026846827.1 | *Drosophila persimilis* |
| Apolygus_lucorum | KAF6213226.1 | *Apolygus lucorum* |
| Drosophila_suzukii | XP_036673287.1 | *Drosophila suzukii* |
| Aphis_glycines | KAE9538112.1 | *Aphis glycines* |
| Drosophila_miranda | XP_017140812.1 | *Drosophila miranda* |
| Lucilia_cuprina | XP_023302899.1 | *Lucilia cuprina* |
| Neodiprion_lecontei | XP_015524986.1 | *Neodiprion lecontei* |
| Fopius_arisanus | XP_011300712.1 | *Fopius arisanus* |
| Myzus_persicae | XP_022173579.1 | *Myzus persicae* |
| Drosophila_obscura | XP_022215202.1 | *Drosophila obscura* |
| Drosophila_novamexicana | XP_030563926.1 | *Drosophila novamexicana* |
| Drosophila_guanche | XP_034131029.1 | *Drosophila guanche* |
| Drosophila_subobscura | XP_034663794.1 | *Drosophila subobscura* |
| Drosophila_busckii | XP_017848011.1 | *Drosophila busckii* |
| Drosophila_virilis | XP_032294873.1 | *Drosophila virilis* |
| Drosophila_willistoni | XP_002073556.1 | *Drosophila willistoni* |
| Drosophila_elegans | XP_017114009.1 | *Drosophila elegans* |
| Drosophila_pseudoobscura | XP_001358796.2 | *Drosophila pseudoobscura* |
| Drosophila_albomicans | XP_034117449.1 | *Drosophila albomicans* |
| Drosophila_takahashii | XP_017015913.1 | *Drosophila takahashii* |
| Drosophila_arizonae | XP_017855619.1 | *Drosophila arizonae* |
| Drosophila_hydei | XP_023175746.2 | *Drosophila hydei* |
| Drosophila_innubila | XP_034488184.1 | *Drosophila innubila* |
| Drosophila_mojavensis | XP_001998760.1 | *Drosophila mojavensis* |
| Stomoxys_calcitrans | XP_013113930.1 | *Stomoxys calcitrans* |
| Drosophila_navojoa | XP_017968702.1 | *Drosophila navojoa* |
| Drosophila_rhopaloa | XP_016978536.1 | *Drosophila rhopaloa* |
| Musca_domestica | XP_005176054.1 | *Musca domestica* |
| Lamprigera_yunnana | KAF5279666.1 | *Lamprigera yunnana* |
| Leptidea_sinapis | VVC98935.1 | *Leptidea sinapis* |
| Aphidius_gifuensis | KAF7993433.1 | *Aphidius gifuensis* |
| Drosophila_grimshawi | XP_001990001.1 | *Drosophila grimshawi* |
| Glossina_fuscipes | XP_037896233.1 | *Glossina fuscipes* |
| Lasius_niger | KMQ92462.1 | *Lasius niger* |
| Clunio_marinus | CRL03231.1 | *Clunio marinus* |
| Contarinia_nasturtii | XP_031620649.1 | *Contarinia nasturtii* |
| Anopheles_albimanus | XP_035785032.1 | *Anopheles albimanus* |
| Anopheles_darlingi | ETN66281.1 | *Anopheles darlingi* |
| Habropoda_laboriosa | XP_017798971.1 | *Habropoda laboriosa* |
| Locusta_migratoria | JAMg_model_8363.1 | *Locusta migratoria* |
| Diabrotica_virgifera_virgifera | XP_028143224.1 | *Diabrotica virgifera virgifera* |
| Operophtera_brumata | KOB68574.1 | *Operophtera brumata* |

**Table S4 List of tree node label, species and accession number of METTL14 gene in insects**

| **Tree node label** | **Accession number** | **Species** |
| --- | --- | --- |
| Bombyx_mandarina | XP_028029337.1 | *Bombyx mandarina* |
| Bombyx_mori | XP_004924405.2 | *Bombyx mori* |
| Spodoptera_exigua | KAF9414146.1 | *Spodoptera exigua* |
| Heliothis_virescens | PCG69680.1, PCG81004.1 | *Heliothis virescens* |
| Helicoverpa_armigera | XP_021186130.1 | *Helicoverpa armigera* |
| Manduca_sexta | XP_030022591.1 | *Manduca sexta* |
| Spodoptera_litura | XP_022823158.1 | *Spodoptera litura* |
| Trichoplusia_ni | XP_026733132.1 | *Trichoplusia ni* |
| Spodoptera_frugiperda | XP_035443576.1, XP_035443990.1 | *Spodoptera frugiperda* |
| Arctia_plantaginis | CAB3234419.1 | *Arctia plantaginis* |
| Ostrinia_furnacalis | XP_028158720.1 | *Ostrinia furnacalis* |
| Galleria_mellonella | XP_026756183.1 | *Galleria mellonella* |
| Amyelois_transitella | XP_013195550.1 | *Amyelois transitella* |
| Vanessa_tameamea | XP_026488012.1 | *Vanessa tameamea* |
| Hyposmocoma_kahamanoa | XP_026322749.1 | *Hyposmocoma kahamanoa* |
| Pieris_rapae | XP_022115997.1 | *Pieris rapae* |
| Aphantopus_hyperantus | XP_034834880.1 | *Aphantopus hyperantus* |
| Plutella_xylostella | XP_011551280.2 | *Plutella xylostella* |
| Chilo_suppressalis | RVE48003.1 | *Chilo suppressalis* |
| Papilio_xuthus | KPI99768.1 | *Papilio xuthus* |
| Danaus_plexippus_plexippus | XP_032518889.1 | *Danaus plexippus plexippus* |
| Eumeta_japonica | GBP49494.1 | *Eumeta japonica* |
| Papilio_polytes | XP_013148094.1 | *Papilio polytes* |
| Bicyclus_anynana | XP_023944713.1 | *Bicyclus anynana* |
| Agrilus_planipennis | XP_018320387.1, XP_025829212.1 | *Agrilus planipennis* |
| Cryptotermes_secundus | XP_023713471.1, XP_033610421.1 | *Cryptotermes secundus* |
| Photinus_pyralis | XP_031347183.1 | *Photinus pyralis* |
| Ctenocephalides_felis | XP_026473557.1 | *Ctenocephalides felis* |
| Thrips_palmi | XP_034229964.1 | *Thrips palmi* |
| Chelonus_insularis | XP_034939255.1 | *Chelonus insularis* |
| Bombus_bifarius | XP_033309867.1 | *Bombus bifarius* |
| Bombus_vancouverensis_nearcticus | XP_033202866.1 | *Bombus vancouverensis nearcticus* |
| Bombus_vosnesenskii | XP_033357485.1 | *Bombus vosnesenskii* |
| Ignelater_luminosus | KAF2899800.1 | *Ignelater luminosus* |
| Frankliniella_occidentalis | KAE8741679.1 | *Frankliniella occidentalis* |
| Bombus_impatiens | XP_003486175.1 | *Bombus impatiens* |
| Bombus_terrestris | XP_003402816.1 | *Bombus terrestris* |
| Habropoda_laboriosa | XP_017789384.1, XP_017789313.1 | *Habropoda laboriosa* |
| Nomia_melanderi | XP_031849658.1 | *Nomia melanderi* |
| Apis_dorsata | XP_006622019.1 | *Apis dorsata* |
| Frieseomelitta_varia | KAF3422349.1 | *Frieseomelitta varia* |
| Apis_cerana | XP_016903943.1 | *Apis cerana* |
| Apis_florea | XP_003698676.1 | *Apis florea* |
| Anopheles_sinensis | KFB49453.1 | *Anopheles sinensis* |
| Apis_mellifera | XP_393391.1 | *Apis mellifera* |
| Megalopta_genalis | XP_033321648.1 | *Megalopta genalis* |
| Ephemera_danica | KAF4522634.1 | *Ephemera danica* |
| Halyomorpha_halys | XP_014283040.1 | *Halyomorpha halys* |
| Ceratina_calcarata | XP_017885386.1 | *Ceratina calcarata* |
| Aedes_albopictus | XP_019535706.1, XP_029709021.1 | *Aedes albopictus* |
| Aedes_aegypti | XP_021702143.1 | *Aedes aegypti* |
| Osmia_bicornis_bicornis | XP_029046277.1 | *Osmia bicornis bicornis* |
| Cloeon_dipterum | CAB3365275.1, CAB3364163.1 | *Cloeon dipterum* |
| Vespa_mandarinia | XP_035739477.1 | *Vespa mandarinia* |
| Vespula_germanica | KAF7390089.1 | *Vespula germanica* |
| Vespula_pensylvanica | KAF7413055.1 | *Vespula pensylvanica* |
| Vespula_vulgaris | KAF7388010.1 | *Vespula vulgaris* |
| Athalia_rosae | XP_012263315.1 | *Athalia rosae* |
| Asbolus_verrucosus | RZC40328.1 | *Asbolus verrucosus* |
| Eufriesea_mexicana | XP_017757340.1 | *Eufriesea mexicana* |
| Microplitis_demolitor | XP_008547285.1 | *Microplitis demolitor* |
| Polistes_canadensis | XP_014602915.1 | *Polistes canadensis* |
| Pediculus_humanus_corporis | XP_002431431.1 | *Pediculus humanus corporis* |
| Anopheles_gambiae | XP_317346.3 | *Anopheles gambiae* |
| Aethina_tumida | XP_019868745.1 | *Aethina tumida* |
| Ooceraea_biroi | XP_011345749.1 | *Ooceraea biroi* |
| Polistes_dominula | XP_015174990.1 | *Polistes dominula* |
| Acromyrmex_echinatior | XP_011059005.1 | *Acromyrmex echinatior* |
| Atta_cephalotes | XP_012056825.1 | *Atta cephalotes* |
| Atta_colombica | XP_018051651.1 | *Atta colombica* |
| Oryctes_borbonicus | KRT84628.1 | *Oryctes borbonicus* |
| Rhynchophorus_ferrugineus | KAF7265822.1 | *Rhynchophorus ferrugineus* |
| Osmia_lignaria | XP_034194464.1 | *Osmia lignaria* |
| Megachile_rotundata | XP_003708161.2 | *Megachile rotundata* |
| Diachasma_alloeum | XP_015113906.1 | *Diachasma alloeum* |
| Trachymyrmex_zeteki | XP_018318018.1 | *Trachymyrmex zeteki* |
| Trachymyrmex_cornetzi | XP_018368258.1 | *Trachymyrmex cornetzi* |
| Temnothorax_curvispinosus | XP_024889385.1 | *Temnothorax curvispinosus* |
| Vollenhovia_emeryi | XP_011861701.1 | *Vollenhovia emeryi* |
| Ceratosolen_solmsi_marchali | XP_011505937.1 | *Ceratosolen solmsi marchali* |
| Dinoponera_quadriceps | XP_014482702.1 | *Dinoponera quadriceps* |
| Apis_cerana_cerana | PBC30532.1 | *Apis cerana cerana* |
| Culex_quinquefasciatus | XP_001867036.1 | *Culex quinquefasciatus* |
| Trachymyrmex_septentrionalis | XP_018338350.1 | *Trachymyrmex septentrionalis* |
| Anoplophora_glabripennis | XP_018561874.1 | *Anoplophora glabripennis* |
| Linepithema_humile | XP_012218973.1 | *Linepithema humile* |
| Pseudomyrmex_gracilis | XP_020287781.1 | *Pseudomyrmex gracilis* |
| Trichomalopsis_sarcophagae | OXU29304.1 | *Trichomalopsis sarcophagae* |
| Cimex_lectularius | XP_014246832.1 | *Cimex lectularius* |
| Leptinotarsa_decemlineata | XP_023029362.1 | *Leptinotarsa decemlineata* |
| Wasmannia_auropunctata | XP_011695743.1 | *Wasmannia auropunctata* |
| Camponotus_floridanus | XP_025264258.1 | *Camponotus floridanus* |
| Neodiprion_lecontei | XP_015514463.1 | *Neodiprion lecontei* |
| Tribolium_castaneum | XP_974982.1 | *Tribolium castaneum* |
| Fopius_arisanus | XP_011304517.1 | *Fopius arisanus* |
| Cyphomyrmex_costatus | XP_018406675.1 | *Cyphomyrmex costatus* |
| Odontomachus_brunneus | XP_032666656.1 | *Odontomachus brunneus* |
| Nylanderia_fulva | XP_029155701.1 | *Nylanderia fulva* |
| Contarinia_nasturtii | XP_031634907.1 | *Contarinia nasturtii* |
| Lasius_niger | KMQ95307.1 | *Lasius niger* |
| Nasonia_vitripennis | XP_016837467.1 | *Nasonia vitripennis* |
| Diaphorina_citri | XP_008479813.1 | *Diaphorina citri* |
| Bemisia_tabaci | XP_018909987.1 | *Bemisia tabaci* |
| Anopheles_stephensi | XP_035915801.1 | *Anopheles stephensi* |
| Coptotermes_formosanus | GFG29635.1, GFG29009.1 | *Coptotermes formosanus* |
| Formica_exsecta | XP_029662651.1 | *Formica exsecta* |
| Copidosoma_floridanum | XP_014206981.1 | *Copidosoma floridanum* |
| Solenopsis_invicta | XP_011159923.1 | *Solenopsis invicta* |
| Dufourea_novaeangliae | XP_015437116.1 | *Dufourea novaeangliae* |
| Aphidius_gifuensis | KAF7997300.1 | *Aphidius gifuensis* |
| Papilio_machaon | XP_014367924.1 | *Papilio machaon* |
| Anopheles_darlingi | ETN58884.1 | *Anopheles darlingi* |
| Sitophilus_oryzae | XP_030754275.1 | *Sitophilus oryzae* |
| Diabrotica_virgifera_virgifera | XP_028130632.1 | *Diabrotica virgifera virgifera* |
| Dendroctonus_ponderosae | XP_019766864.1 | *Dendroctonus ponderosae* |
| Anopheles_albimanus | XP_035776345.1 | *Anopheles albimanus* |
| Monomorium_pharaonis | XP_012533546.1 | *Monomorium pharaonis* |
| Harpegnathos_saltator | XP_019695918.1 | *Harpegnathos saltator* |
| Pogonomyrmex_barbatus | XP_011636458.1 | *Pogonomyrmex barbatus* |
| Drosophila_biarmipes | XP_016967622.1 | *Drosophila biarmipes* |
| Belonocnema_treatae | XP_033228574.1 | *Belonocnema treatae* |
| Cephus_cinctus | XP_015604153.1 | *Cephus cinctus* |
| Zootermopsis_nevadensis | XP_021922333.1 | *Zootermopsis nevadensis* |
| Callosobruchus_maculatus | VEN63447.1 | *Callosobruchus maculatus* |
| Hermetia_illucens | XP_037903057.1 | *Hermetia illucens* |
| Ceratitis_capitata | CAD7000340.1 | *Ceratitis capitata* |
| Trichogramma_pretiosum | XP_014223352.1 | *Trichogramma pretiosum* |
| Drosophila_guanche | XP_034125006.1 | *Drosophila guanche* |
| Drosophila_obscura | XP_022225558.1 | *Drosophila obscura* |
| Drosophila_subobscura | XP_034670304.1 | *Drosophila subobscura* |
| Sarcophaga_bullata | TMW53226.1 | *Sarcophaga bullata* |
| Drosophila_willistoni | XP_002064446.1 | *Drosophila willistoni* |
| Onthophagus_taurus | XP_022909160.1 | *Onthophagus taurus* |
| Drosophila_miranda | XP_017151606.1 | *Drosophila miranda* |
| Drosophila_persimilis | XP_002014836.1 | *Drosophila persimilis* |
| Drosophila_pseudoobscura | XP_001356597.3 | *Drosophila pseudoobscura* |
| Drosophila_subpulchrella | XP_037711280.1 | *Drosophila subpulchrella* |
| Drosophila_suzukii | XP_016924178.1 | *Drosophila suzukii* |
| Drosophila_yakuba | XP_002088776.1 | *Drosophila yakuba* |
| Zeugodacus_cucurbitae | XP_011180293.1 | *Zeugodacus cucurbitae* |
| Drosophila_melanogaster | NP_609205.1 | *Drosophila melanogaster* |
| Drosophila_ficusphila | XP_017047968.1 | *Drosophila ficusphila* |
| Drosophila_sechellia | XP_002036189.1 | *Drosophila sechellia* |
| Rhagoletis_pomonella | XP_036328428.1 | *Rhagoletis pomonella* |
| Drosophila_elegans | XP_017132657.1 | *Drosophila elegans* |
| Drosophila_rhopaloa | XP_016989355.1 | *Drosophila rhopaloa* |
| Drosophila_simulans | XP_002078624.1 | *Drosophila simulans* |
| Apolygus_lucorum | KAF6213089.1 | *Apolygus lucorum* |
| Musca_domestica | XP_005187763.1 | *Musca domestica* |
| Teleopsis_dalmanni | XP_037957915.1 | *Teleopsis dalmanni* |
| Rhagoletis_zephyria | XP_017480566.1, XP_017489009.1 | *Rhagoletis zephyria* |
| Drosophila_eugracilis | XP_017068613.1 | *Drosophila eugracilis* |
| Drosophila_mauritiana | XP_033155733.1 | *Drosophila mauritiana* |
| Bactrocera_latifrons | XP_018796376.1 | *Bactrocera latifrons* |
| Drosophila_erecta | XP_001970274.1 | *Drosophila erecta* |
| Drosophila_kikkawai | XP_017029555.1 | *Drosophila kikkawai* |
| Drosophila_serrata | XP_020799005.1 | *Drosophila serrata* |
| Drosophila_bipectinata | XP_017097828.1 | *Drosophila bipectinata* |
| Lamprigera_yunnana | KAF5277462.1 | *Lamprigera yunnana* |
| Bactrocera_dorsalis | XP_011199013.1 | *Bactrocera dorsalis* |
| Drosophila_takahashii | XP_017016124.1 | *Drosophila takahashii* |
| Stomoxys_calcitrans | XP_013105928.1 | *Stomoxys calcitrans* |
| Scaptodrosophila_lebanonensis | XP_030383882.1 | *Scaptodrosophila lebanonensis* |
| Drosophila_ananassae | XP_001962040.1 | *Drosophila ananassae* |
| Nicrophorus_vespilloides | XP_017773393.1 | *Nicrophorus vespilloides* |
| Bactrocera_oleae | XP_014087401.1, XP_014088588.1 | *Bactrocera oleae* |
| Drosophila_innubila | XP_034474151.1 | *Drosophila innubila* |
| Temnothorax_longispinosus | TGZ51408.1 | *Temnothorax longispinosus* |
| Drosophila_novamexicana | XP_030556800.1 | *Drosophila novamexicana* |
| Drosophila_virilis | XP_002052643.1 | *Drosophila virilis* |
| Glossina_fuscipes | XP_037883028.1 | *Glossina fuscipes* |
| Lucilia_sericata | XP_037820969.1 | *Lucilia sericata* |
| Drosophila_navojoa | XP_017954488.1 | *Drosophila navojoa* |
| Lucilia_cuprina | XP_023304760.1 | *Lucilia cuprina* |
| Drosophila_arizonae | XP_017859921.1 | *Drosophila arizonae* |
| Drosophila_mojavensis | XP_002002031.1 | *Drosophila mojavensis* |
| Myzus_persicae | XP_022169676.1 | *Myzus persicae* |
| Drosophila_hydei | XP_023166108.1 | *Drosophila hydei* |
| Drosophila_grimshawi | XP_001989197.2 | *Drosophila grimshawi* |
| Drosophila_albomicans | XP_034101378.1 | *Drosophila albomicans* |
| Acyrthosiphon_pisum | XP_001948323.1, XP_001950099.1 | *Acyrthosiphon pisum* |
| Aphis_gossypii | XP_027838799.1, XP_027849411.1 | *Aphis gossypii* |
| Aphis_craccivora | KAF0771529.1, KAF0756185.1 | *Aphis craccivora* |
| Melanaphis_sacchari | XP_025195339.1, XP_025208649.1 | *Melanaphis sacchari* |
| Diuraphis_noxia | XP_015379656.1 | *Diuraphis noxia* |
| Rhopalosiphum_maidis | XP_026805082.1 | *Rhopalosiphum maidis* |
| Clunio_marinus | CRK86719.1 | *Clunio marinus* |
| Sipha_flava | XP_025423473.1 | *Sipha flava* |
| Cinara_cedri | VVC24649.1 | *Cinara cedri* |
| Drosophila_busckii | XP_017834825.1 | *Drosophila busckii* |
| Abscondita_terminalis | KAF5296451.1 | *Abscondita terminalis* |
| Melipona_quadrifasciata | KOX80044.1 | *Melipona quadrifasciata* |
| Orussus_abietinus | XP_012282380.1 | *Orussus abietinus* |
| Aphis_glycines | KAE9534352.1, KAE9522259.1 | *Aphis glycines* |
| Leptidea_sinapis | VVC94570.1 | *Leptidea sinapis* |
| Nesidiocoris_tenuis | CAB0003557.1 | *Nesidiocoris tenuis* |
| Locusta_migratoria | JAMg_model_2092.1 | *Locusta migratoria* |
| Blattella_germanica | PSN50207.1 | *Blattella germanica* |

**Table S5 List of tree node label, species and accession number of YTHDC gene in insects**

| **Tree node label** | **Accession number** | **Species** |
| --- | --- | --- |
| Bombyx_mori | XP_004931496.2 | *Bombyx mori* |
| Bombyx_mandarina | XP_028031043.1 | *Bombyx mandarina* |
| Arctia_plantaginis | CAB3259921.1 | *Arctia plantaginis* |
| Amyelois_transitella | XP_013190567.1 | *Amyelois transitella* |
| Chilo_suppressalis | RVE41834.1 | *Chilo suppressalis* |
| Galleria_mellonella | XP_026762523.1 | *Galleria mellonella* |
| Trichoplusia_ni | XP_026730454.1 | *Trichoplusia ni* |
| Spodoptera_litura | XP_022835342.1 | *Spodoptera litura* |
| Spodoptera_frugiperda | XP_035445160.1 | *Spodoptera frugiperda* |
| Ostrinia_furnacalis | XP_028175992.1 | *Ostrinia furnacalis* |
| Hyposmocoma_kahamanoa | XP_026316753.1 | *Hyposmocoma kahamanoa* |
| Danaus_plexippus_plexippus | XP_032520739.1 | *Danaus plexippus plexippus* |
| Heliothis_virescens | PCG72237.1, PCG72245.1 | *Heliothis virescens* |
| Aphantopus_hyperantus | XP_034831521.1 | *Aphantopus hyperantus* |
| Papilio_machaon | XP_014368863.1 | *Papilio machaon* |
| Plutella_xylostella | XP_037974788.1 | *Plutella xylostella* |
| Papilio_polytes | XP_013146725.1 | *Papilio polytes* |
| Pieris_rapae | XP_022113565.1 | *Pieris rapae* |
| Helicoverpa_armigera | XP_021181528.1 | *Helicoverpa armigera* |
| Eumeta_japonica | GBP19843.1 | *Eumeta japonica* |
| Bicyclus_anynana | XP_023936770.1 | *Bicyclus anynana* |
| Manduca_sexta | XP_030036110.1 | *Manduca sexta* |
| Leptidea_sinapis | VVC86746.1 | *Leptidea sinapis* |
| Vanessa_tameamea | XP_026499160.1 | *Vanessa tameamea* |
| Operophtera_brumata | KOB78084.1 | *Operophtera brumata* |
| Papilio_xuthus | KPI97362.1 | *Papilio xuthus* |
| Cryptotermes_secundus | XP_023713379.1, XP_023712706.1 | *Cryptotermes secundus* |
| Zootermopsis_nevadensis | XP_021929385.1, XP_021937348.1 | *Zootermopsis nevadensis* |
| Ignelater_luminosus | KAF2902905.1 | *Ignelater luminosus* |
| Photinus_pyralis | XP_031347580.1 | *Photinus pyralis* |
| Onthophagus_taurus | XP_022911608.1 | *Onthophagus taurus* |
| Abscondita_terminalis | KAF5280547.1 | *Abscondita terminalis* |
| Anoplophora_glabripennis | XP_018577954.1 | *Anoplophora glabripennis* |
| Agrilus_planipennis | XP_018323290.1, XP_025836712.1 | *Agrilus planipennis* |
| Tribolium_castaneum | XP_969804.2 | *Tribolium castaneum* |
| Blattella_germanica | PSN37472.1 | *Blattella germanica* |
| Rhynchophorus_ferrugineus | KAF7285969.1 | *Rhynchophorus ferrugineus* |
| Asbolus_verrucosus | RZC40283.1 | *Asbolus verrucosus* |
| Ctenocephalides_felis | XP_026480156.1, XP_026480114.1, XP_026480049.1 | *Ctenocephalides felis* |
| Neodiprion_lecontei | XP_015516731.1 | *Neodiprion lecontei* |
| Locusta_migratoria | JAMg_model_14479.1 | *Locusta migratoria* |
| Fopius_arisanus | XP_011299102.1 | *Fopius arisanus* |
| Diabrotica_virgifera_virgifera | XP_028149969.1 | *Diabrotica virgifera virgifera* |
| Frankliniella_occidentalis | KAE8740230.1 | *Frankliniella occidentalis* |
| Leptinotarsa_decemlineata | XP_023016598.1 | *Leptinotarsa decemlineata* |
| Nicrophorus_vespilloides | XP_017768326.1 | *Nicrophorus vespilloides* |
| Athalia_rosae | XP_012263380.1 | *Athalia rosae* |
| Cephus_cinctus | XP_015587583.1 | *Cephus cinctus* |
| Osmia_bicornis_bicornis | XP_029050552.1 | *Osmia bicornis bicornis* |
| Osmia_lignaria | XP_034185805.1 | *Osmia lignaria* |
| Eufriesea_mexicana | XP_017767385.1 | *Eufriesea mexicana* |
| Diachasma_alloeum | XP_015118637.1 | *Diachasma alloeum* |
| Bombus_terrestris | XP_003398194.1 | *Bombus terrestris* |
| Polistes_dominula | XP_015171044.1 | *Polistes dominula* |
| Apis_cerana | XP_016903802.1 | *Apis cerana* |
| Habropoda_laboriosa | XP_017789685.1 | *Habropoda laboriosa* |
| Melipona_quadrifasciata | KOX78177.1 | *Melipona quadrifasciata* |
| Apis_dorsata | XP_006613940.1 | *Apis dorsata* |
| Vespula_germanica | KAF7399625.1 | *Vespula germanica* |
| Apis_mellifera | XP_395221.4 | *Apis mellifera* |
| Vespula_pensylvanica | KAF7423658.1 | *Vespula pensylvanica* |
| Vespula_vulgaris | KAF7396604.1 | *Vespula vulgaris* |
| Bombus_bifarius | XP_033299410.1 | *Bombus bifarius* |
| Bombus_impatiens | XP_012240497.1 | *Bombus impatiens* |
| Bombus_vancouverensis_nearcticus | XP_033202336.1 | *Bombus vancouverensis nearcticus* |
| Bombus_vosnesenskii | XP_033343380.1 | *Bombus vosnesenskii* |
| Frieseomelitta_varia | KAF3420202.1 | *Frieseomelitta varia* |
| Nomia_melanderi | XP_031825916.1 | *Nomia melanderi* |
| Apis_cerana_cerana | PBC29594.1 | *Apis cerana cerana* |
| Odontomachus_brunneus | XP_032663144.1 | *Odontomachus brunneus* |
| Wasmannia_auropunctata | XP_011695246.1 | *Wasmannia auropunctata* |
| Megalopta_genalis | XP_033325468.1 | *Megalopta genalis* |
| Linepithema_humile | XP_012225148.1 | *Linepithema humile* |
| Apis_florea | XP_003698309.1 | *Apis florea* |
| Dufourea_novaeangliae | XP_015439599.1 | *Dufourea novaeangliae* |
| Vespa_mandarinia | XP_035718955.1 | *Vespa mandarinia* |
| Solenopsis_invicta | XP_011168938.1 | *Solenopsis invicta* |
| Cyphomyrmex_costatus | XP_018399392.1 | *Cyphomyrmex costatus* |
| Microplitis_demolitor | XP_008561185.1 | *Microplitis demolitor* |
| Ooceraea_biroi | XP_011337952.1 | *Ooceraea biroi* |
| Ceratina_calcarata | XP_026667650.1 | *Ceratina calcarata* |
| Thrips_palmi | XP_034249974.1 | *Thrips palmi* |
| Temnothorax_curvispinosus | XP_024881572.1 | *Temnothorax curvispinosus* |
| Vollenhovia_emeryi | XP_011866786.1 | *Vollenhovia emeryi* |
| Chelonus_insularis | XP_034946843.1 | *Chelonus insularis* |
| Monomorium_pharaonis | XP_012536917.1 | *Monomorium pharaonis* |
| Nylanderia_fulva | XP_029174627.1 | *Nylanderia fulva* |
| Sitophilus_oryzae | XP_030760572.1 | *Sitophilus oryzae* |
| Camponotus_floridanus | XP_011253687.1 | *Camponotus floridanus* |
| Pseudomyrmex_gracilis | XP_020282017.1 | *Pseudomyrmex gracilis* |
| Pogonomyrmex_barbatus | XP_011634510.1 | *Pogonomyrmex barbatus* |
| Trachymyrmex_zeteki | XP_018305074.1 | *Trachymyrmex zeteki* |
| Atta_cephalotes | XP_012058638.1 | *Atta cephalotes* |
| Atta_colombica | XP_018043733.1 | *Atta colombica* |
| Trachymyrmex_septentrionalis | XP_018343111.1 | *Trachymyrmex septentrionalis* |
| Dendroctonus_ponderosae | XP_019755440.1 | *Dendroctonus ponderosae* |
| Orussus_abietinus | XP_012274238.1 | *Orussus abietinus* |
| Harpegnathos_saltator | XP_011154191.1 | *Harpegnathos saltator* |
| Acromyrmex_echinatior | XP_011054414.1 | *Acromyrmex echinatior* |
| Dinoponera_quadriceps | XP_014474127.1 | *Dinoponera quadriceps* |
| Belonocnema_treatae | XP_033209600.1 | *Belonocnema treatae* |
| Trachymyrmex_cornetzi | XP_018376609.1 | *Trachymyrmex cornetzi* |
| Formica_exsecta | XP_029676764.1 | *Formica exsecta* |
| Aphidius_gifuensis | KAF7992969.1 | *Aphidius gifuensis* |
| Ceratosolen_solmsi_marchali | XP_011499020.1 | *Ceratosolen solmsi marchali* |
| Temnothorax_longispinosus | TGZ49014.1 | *Temnothorax longispinosus* |
| Copidosoma_floridanum | XP_014215728.1 | *Copidosoma floridanum* |
| Aethina_tumida | XP_019866243.1 | *Aethina tumida* |
| Nasonia_vitripennis | XP_001604858.2 | *Nasonia vitripennis* |
| Hermetia_illucens | XP_037905739.1 | *Hermetia illucens* |
| Trichogramma_pretiosum | XP_014228483.1 | *Trichogramma pretiosum* |
| Lasius_niger | KMQ96841.1 | *Lasius niger* |
| Bemisia_tabaci | XP_018901813.1 | *Bemisia tabaci* |
| Culex_quinquefasciatus | XP_001849154.1 | *Culex quinquefasciatus* |
| Contarinia_nasturtii | XP_031634606.1 | *Contarinia nasturtii* |
| Aedes_albopictus | XP_019933141.1 | *Aedes albopictus* |
| Aedes_aegypti | XP_001661159.1 | *Aedes aegypti* |
| Pediculus_humanus_corporis | XP_002427239.1 | *Pediculus humanus corporis* |
| Trichomalopsis_sarcophagae | OXU19416.1 | *Trichomalopsis sarcophagae* |
| Megachile_rotundata | XP_003708670.2 | *Megachile rotundata* |
| Trichogramma_brassicae | CAB0029133.1 | *Trichogramma brassicae* |
| Callosobruchus_maculatus | VEN44205.1 | *Callosobruchus maculatus* |
| Cloeon_dipterum | CAB3374621.1 | *Cloeon dipterum* |
| Anopheles_darlingi | ETN64938.1 | *Anopheles darlingi* |
| Clunio_marinus | CRK99451.1 | *Clunio marinus* |
| Anopheles_stephensi | XP_035914346.1 | *Anopheles stephensi* |
| Anopheles_albimanus | XP_035790890.1 | *Anopheles albimanus* |
| Drosophila_sechellia | XP_032573220.1 | *Drosophila sechellia* |
| Drosophila_erecta | XP_026832853.1 | *Drosophila erecta* |
| Drosophila_rhopaloa | XP_016975531.1 | *Drosophila rhopaloa* |
| Drosophila_ficusphila | XP_017058042.1 | *Drosophila ficusphila* |
| Scaptodrosophila_lebanonensis | XP_030378134.1 | *Scaptodrosophila lebanonensis* |
| Drosophila_serrata | XP_020811705.1 | *Drosophila serrata* |
| Drosophila_subpulchrella | XP_037720181.1 | *Drosophila subpulchrella* |
| Drosophila_mauritiana | XP_033158309.1 | *Drosophila mauritiana* |
| Drosophila_simulans | XP_016030234.1 | *Drosophila simulans* |
| Drosophila_suzukii | XP_036670453.1 | *Drosophila suzukii* |
| Drosophila_takahashii | XP_017011424.1 | *Drosophila takahashii* |
| Anopheles_sinensis | KFB47137.1 | *Anopheles sinensis* |
| Drosophila_biarmipes | XP_016966523.1 | *Drosophila biarmipes* |
| Drosophila_yakuba | XP_002093557.1 | *Drosophila yakuba* |
| Drosophila_kikkawai | XP_017021329.1 | *Drosophila kikkawai* |
| Drosophila_melanogaster | NP_647811.2 | *Drosophila melanogaster* |
| Drosophila_novamexicana | XP_030572459.1 | *Drosophila novamexicana* |
| Anopheles_gambiae | XP_001688755.1 | *Anopheles gambiae* |
| Drosophila_eugracilis | XP_017071429.1 | *Drosophila eugracilis* |
| Drosophila_virilis | XP_002046838.2 | *Drosophila virilis* |
| Drosophila_navojoa | XP_017955029.1 | *Drosophila navojoa* |
| Drosophila_arizonae | XP_017864073.1 | *Drosophila arizonae* |
| Drosophila_mojavensis | XP_002007461.2 | *Drosophila mojavensis* |
| Drosophila_guanche | XP_034136733.1 | *Drosophila guanche* |
| Drosophila_subobscura | XP_034658093.1 | *Drosophila subobscura* |
| Drosophila_obscura | XP_022225017.1 | *Drosophila obscura* |
| Drosophila_hydei | XP_023173281.1 | *Drosophila hydei* |
| Drosophila_elegans | XP_017127640.1 | *Drosophila elegans* |
| Drosophila_innubila | XP_034481636.1 | *Drosophila innubila* |
| Teleopsis_dalmanni | XP_037941807.1 | *Teleopsis dalmanni* |
| Drosophila_ananassae | XP_001957138.1 | *Drosophila ananassae* |
| Drosophila_busckii | XP_017834951.1 | *Drosophila busckii* |
| Drosophila_bipectinata | XP_017098472.1 | *Drosophila bipectinata* |
| Bactrocera_dorsalis | XP_011205865.1 | *Bactrocera dorsalis* |
| Bactrocera_latifrons | XP_018787966.1 | *Bactrocera latifrons* |
| Cimex_lectularius | XP_014248285.1 | *Cimex lectularius* |
| Sarcophaga_bullata | TMW42074.1 | *Sarcophaga bullata* |
| Drosophila_willistoni | XP_002068514.3 | *Drosophila willistoni* |
| Drosophila_miranda | XP_017137843.1 | *Drosophila miranda* |
| Drosophila_persimilis | XP_026847532.1 | *Drosophila persimilis* |
| Drosophila_pseudoobscura | XP_002135520.1 | *Drosophila pseudoobscura* |
| Drosophila_albomicans | XP_034104636.1 | *Drosophila albomicans* |
| Lucilia_cuprina | XP_023292455.1 | *Lucilia cuprina* |
| Lucilia_sericata | XP_037818608.1 | *Lucilia sericata* |
| Glossina_fuscipes | XP_037880691.1 | *Glossina fuscipes* |
| Zeugodacus_cucurbitae | XP_011195560.1 | *Zeugodacus cucurbitae* |
| Stomoxys_calcitrans | XP_013118835.1 | *Stomoxys calcitrans* |
| Ceratitis_capitata | CAD7004698.1 | *Ceratitis capitata* |
| Bactrocera_oleae | XP_014103061.2 | *Bactrocera oleae* |
| Halyomorpha_halys | XP_014289436.1 | *Halyomorpha halys* |
| Rhagoletis_pomonella | XP_036336384.1 | *Rhagoletis pomonella* |
| Rhagoletis_zephyria | XP_017461432.1 | *Rhagoletis zephyria* |
| Musca_domestica | XP_005179046.1 | *Musca domestica* |
| Nesidiocoris_tenuis | CAB0013311.1 | *Nesidiocoris tenuis* |
| Drosophila_grimshawi | XP_001997660.2, XP_001983824.2 | *Drosophila grimshawi* |
| Apolygus_lucorum | KAF6202688.1 | *Apolygus lucorum* |
| Ephemera_danica | KAF4533255.1 | *Ephemera danica* |

**Table S6 List of tree node label, species and accession number of FL2D gene in insects**

| **Tree node label** | **Accession number** | **Species** |
| --- | --- | --- |
| Drosophila_melanogaster | NP_523732.2 | *Drosophila melanogaster* |
| Drosophila_mauritiana | XP_033151885.1 | *Drosophila mauritiana* |
| Drosophila_sechellia | XP_002033752.1 | *Drosophila sechellia* |
| Drosophila_simulans | XP_016027436.1 | *Drosophila simulans* |
| Drosophila_erecta | XP_001975725.1 | *Drosophila erecta* |
| Drosophila_yakuba | XP_002090866.1 | *Drosophila yakuba* |
| Drosophila_subpulchrella | XP_037728388.1 | *Drosophila subpulchrella* |
| Drosophila_suzukii | XP_016943429.1 | *Drosophila suzukii* |
| Drosophila_eugracilis | XP_017069114.1 | *Drosophila eugracilis* |
| Drosophila_biarmipes | XP_016957390.1 | *Drosophila biarmipes* |
| Drosophila_ficusphila | XP_017051190.1 | *Drosophila ficusphila* |
| Drosophila_elegans | XP_017113148.1 | *Drosophila elegans* |
| Drosophila_takahashii | XP_017003361.1 | *Drosophila takahashii* |
| Drosophila_ananassae | XP_032307103.1 | *Drosophila ananassae* |
| Drosophila_bipectinata | XP_017106942.1 | *Drosophila bipectinata* |
| Drosophila_serrata | XP_020815236.1 | *Drosophila serrata* |
| Drosophila_kikkawai | XP_017029739.1 | *Drosophila kikkawai* |
| Drosophila_subobscura | XP_034652880.1 | *Drosophila subobscura* |
| Drosophila_guanche | XP_034136028.1 | *Drosophila guanche* |
| Drosophila_miranda | XP_017149592.1 | *Drosophila miranda* |
| Drosophila_innubila | XP_034479769.1 | *Drosophila innubila* |
| Drosophila_rhopaloa | XP_016979574.1 | *Drosophila rhopaloa* |
| Drosophila_obscura | XP_022220211.1 | *Drosophila obscura* |
| Drosophila_obscura | XP_022232930.1 | *Drosophila obscura* |
| Drosophila_albomicans | XP_034101732.1 | *Drosophila albomicans* |
| Scaptodrosophila_lebanonensis | XP_030384404.1 | *Scaptodrosophila lebanonensis* |
| Drosophila_hydei | XP_023164871.1 | *Drosophila hydei* |
| Drosophila_navojoa | XP_017961208.1 | *Drosophila navojoa* |
| Drosophila_persimilis | XP_002018003.2 | *Drosophila persimilis* |
| Drosophila_pseudoobscura | XP_001361607.3 | *Drosophila pseudoobscura* |
| Drosophila_virilis | XP_032292721.1 | *Drosophila virilis* |
| Drosophila_mojavensis | XP_002006313.1 | *Drosophila mojavensis* |
| Drosophila_grimshawi | XP_001985842.1 | *Drosophila grimshawi* |
| Drosophila_novamexicana | XP_030554129.1 | *Drosophila novamexicana* |
| Drosophila_willistoni | XP_023033750.1 | *Drosophila willistoni* |
| Drosophila_busckii | XP_017839024.1 | *Drosophila busckii* |
| Sarcophaga_bullata | TMW51352.1 | *Sarcophaga bullata* |
| Drosophila_rhopaloa | XP_016985893.1 | *Drosophila rhopaloa* |
| Lucilia_sericata | XP_037819047.1 | *Lucilia sericata* |
| Lucilia_cuprina | XP_023297132.1 | *Lucilia cuprina* |
| Bactrocera_dorsalis | XP_011200969.1 | *Bactrocera dorsalis* |
| Bactrocera_oleae | XP_014088619.1 | *Bactrocera oleae* |
| Zeugodacus_cucurbitae | XP_011189826.1 | *Zeugodacus cucurbitae* |
| Rhagoletis_zephyria | XP_017470920.1 | *Rhagoletis zephyria* |
| Rhagoletis_pomonella | XP_036333575.1 | *Rhagoletis pomonella* |
| Bactrocera_latifrons | XP_018799504.1 | *Bactrocera latifrons* |
| Stomoxys_calcitrans | XP_013119542.1 | *Stomoxys calcitrans* |
| Musca_domestica | XP_005177555.1 | *Musca domestica* |
| Glossina_fuscipes | XP_037887862.1 | *Glossina fuscipes* |
| Teleopsis_dalmanni | XP_037955062.1 | *Teleopsis dalmanni* |
| Teleopsis_dalmanni | XP_037948558.1 | *Teleopsis dalmanni* |
| Hermetia_illucens | XP_037917482.1 | *Hermetia illucens* |
| Culex_quinquefasciatus | XP_001848362.1 | *Culex quinquefasciatus* |
| Aedes_albopictus | XP_019931319.1 | *Aedes albopictus* |
| Aedes_aegypti | XP_001663177.1 | *Aedes aegypti* |
| Contarinia_nasturtii | XP_031622368.1 | *Contarinia nasturtii* |
| Ctenocephalides_felis | XP_026472852.1 | *Ctenocephalides felis* |
| Culex_quinquefasciatus | XP_001868480.1 | *Culex quinquefasciatus* |
| Cephus_cinctus | XP_015599942.1 | *Cephus cinctus* |
| Orussus_abietinus | XP_012270167.1 | *Orussus abietinus* |
| Athalia_rosae | XP_020708876.1 | *Athalia rosae* |
| Anopheles_sinensis | KFB35615.1 | *Anopheles sinensis* |
| Nasonia_vitripennis | XP_003424468.1 | *Nasonia vitripennis* |
| Neodiprion_lecontei | XP_015513129.1 | *Neodiprion lecontei* |
| Anopheles_albimanus | XP_035786748.1 | *Anopheles albimanus* |
| Bombus_bifarius | XP_033314085.1 | *Bombus bifarius* |
| Bombus_impatiens | XP_012249246.1 | *Bombus impatiens* |
| Bombus_terrestris | XP_012169071.1 | *Bombus terrestris* |
| Bombus_vancouverensis_nearcticus | XP_033190210.1 | *Bombus vancouverensis nearcticus* |
| Bombus_vosnesenskii | XP_033348794.1 | *Bombus vosnesenskii* |
| Belonocnema_treatae | XP_033230326.1 | *Belonocnema treatae* |
| Melipona_quadrifasciata | KOX67774.1 | *Melipona quadrifasciata* |
| Anopheles_darlingi | ETN63002.1 | *Anopheles darlingi* |
| Linepithema_humile | XP_012224928.1 | *Linepithema humile* |
| Nylanderia_fulva | XP_029164338.1 | *Nylanderia fulva* |
| Camponotus_floridanus | XP_011258753.1 | *Camponotus floridanus* |
| Atta_cephalotes | XP_012061791.1 | *Atta cephalotes* |
| Atta_colombica | XP_018048902.1 | *Atta colombica* |
| Formica_exsecta | XP_029676355.1 | *Formica exsecta* |
| Vollenhovia_emeryi | XP_011873950.1 | *Vollenhovia emeryi* |
| Apis_dorsata | XP_006615159.1 | *Apis dorsata* |
| Apis_mellifera | XP_006566329.1 | *Apis mellifera* |
| Apis_cerana_cerana | PBC31280.1 | *Apis cerana cerana* |
| Apis_cerana | XP_016914837.1 | *Apis cerana* |
| Monomorium_pharaonis | XP_012525956.1 | *Monomorium pharaonis* |
| Wasmannia_auropunctata | XP_011701827.1 | *Wasmannia auropunctata* |
| Odontomachus_brunneus | XP_032676278.1 | *Odontomachus brunneus* |
| Pogonomyrmex_barbatus | XP_011647104.1 | *Pogonomyrmex barbatus* |
| Aphidius_gifuensis | KAF7993230.1 | *Aphidius gifuensis* |
| Leptinotarsa_decemlineata | XP_023019229.1 | *Leptinotarsa decemlineata* |
| Harpegnathos_saltator | XP_011134848.1 | *Harpegnathos saltator* |
| Callosobruchus_maculatus | VEN45117.1 | *Callosobruchus maculatus* |
| Vespula_germanica | KAF7386550.1 | *Vespula germanica* |
| Vespula_pensylvanica | KAF7407107.1 | *Vespula pensylvanica* |
| Vespula_vulgaris | KAF7384880.1 | *Vespula vulgaris* |
| Anopheles_stephensi | XP_035901105.1 | *Anopheles stephensi* |
| Vespa_mandarinia | XP_035723086.1 | *Vespa mandarinia* |
| Microplitis_demolitor | XP_014299773.1 | *Microplitis demolitor* |
| Pediculus_humanus_corporis | XP_002431432.1 | *Pediculus humanus corporis* |
| Diabrotica_virgifera_virgifera | XP_028145213.1 | *Diabrotica virgifera virgifera* |
| Dinoponera_quadriceps | XP_014468853.1 | *Dinoponera quadriceps* |
| Ephemera_danica | KAF4528646.1 | *Ephemera danica* |
| Nilaparvata_lugens | Sfur05226 | *Nilaparvata lugens* |
| Cotesia_vestalis | Nlug26297 | *Cotesia vestalis* |
| Bemisia_tabaci | XP_018914611.1 | *Bemisia tabaci* |
| Ostrinia_furnacalis | XP_028169814.1 | *Ostrinia furnacalis* |
| Papilio_polytes | XP_013148295.1 | *Papilio polytes* |
| Lamprigera_yunnana | KAF5277460.1 | *Lamprigera yunnana* |
| Ignelater_luminosus | KAF2893495.1 | *Ignelater luminosus* |
| Photinus_pyralis | XP_031353041.1 | *Photinus pyralis* |
| Manduca_sexta | XP_030020275.1 | *Manduca sexta* |
| Plutella_xylostella | XP_011547887.2 | *Plutella xylostella* |
| Chilo_suppressalis | RVE49843.1 | *Chilo suppressalis* |
| Trichomalopsis_sarcophagae | OXU31401.1 | *Trichomalopsis sarcophagae* |
| Hyposmocoma_kahamanoa | XP_026321266.1 | *Hyposmocoma kahamanoa* |
| Cimex_lectularius | XP_014244543.1 | *Cimex lectularius* |
| Cloeon_dipterum | CAB3364017.1 | *Cloeon dipterum* |
| Sipha_flava | XP_025421064.1 | *Sipha flava* |
| Sogatella_furcifera | HaxR_016023 | *Sogatella furcifera* |
| Blattella_germanica | PSN54928.1 | *Blattella germanica* |
| Sipha_flava | XP_025420411.1 | *Sipha flava* |
| Ceratitis_capitata | CAD7012146.1 | *Ceratitis capitata* |
| Diuraphis_noxia | XP_015370775.1 | *Diuraphis noxia* |
| Teleopsis_dalmanni | XP_037954326.1 | *Teleopsis dalmanni* |
| Vespula_pensylvanica | KAF7407108.1 | *Vespula pensylvanica* |
| Vespula_vulgaris | KAF7384881.1 | *Vespula vulgaris* |
| Vespula_germanica | KAF7386551.1 | *Vespula germanica* |
| Papilio_machaon | XP_014371206.1 | *Papilio machaon* |
| Eumeta_japonica | GBP50624.1 | *Eumeta japonica* |

Purified BmMETTL3-His

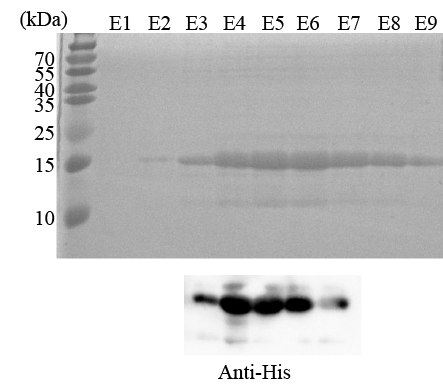

**Figure S1. Expression and purification of BmMETTL3** Recombinant His-tagged *BmMETTL3* protein was expressed by *E. coli*. The *BmMETTL3* was purified by nickel chromatography and subjected to SDS-PAGE through a 10% SDS-polyacrylamide gel, and then stained with CBB. The same samples were analyzed by western blotting with the anti-His antibody. Molecular mass markers are shown to the right of the gel. Samples are as follows: lane 1-9.

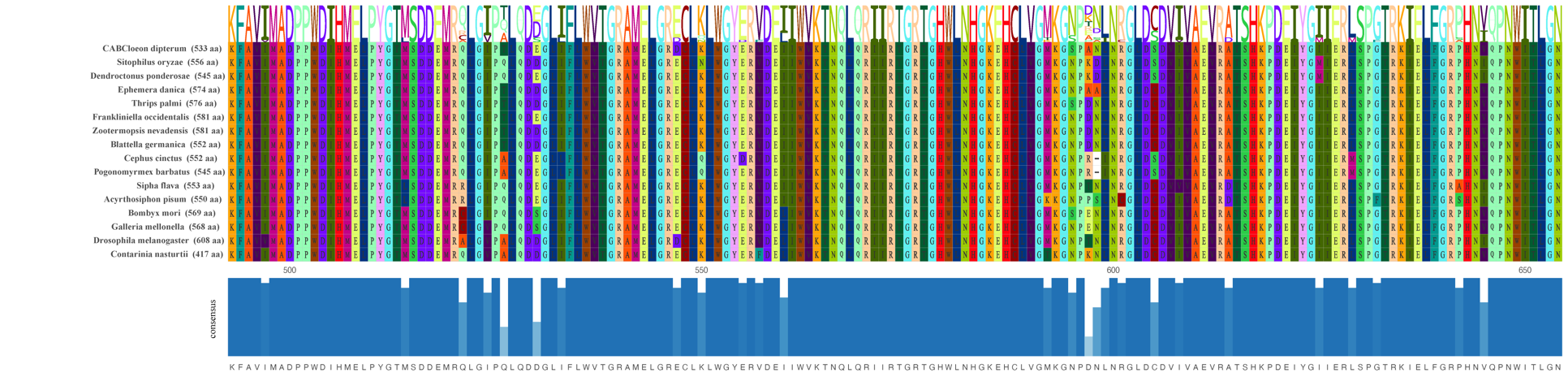

**Figure S2.** Multiple sequence alignments of MT-A70 domain of METTL3 from various species. Alignments were done with CLC Sequence Viewer7. Accession numbers for all sequences are provided in the Table S1.

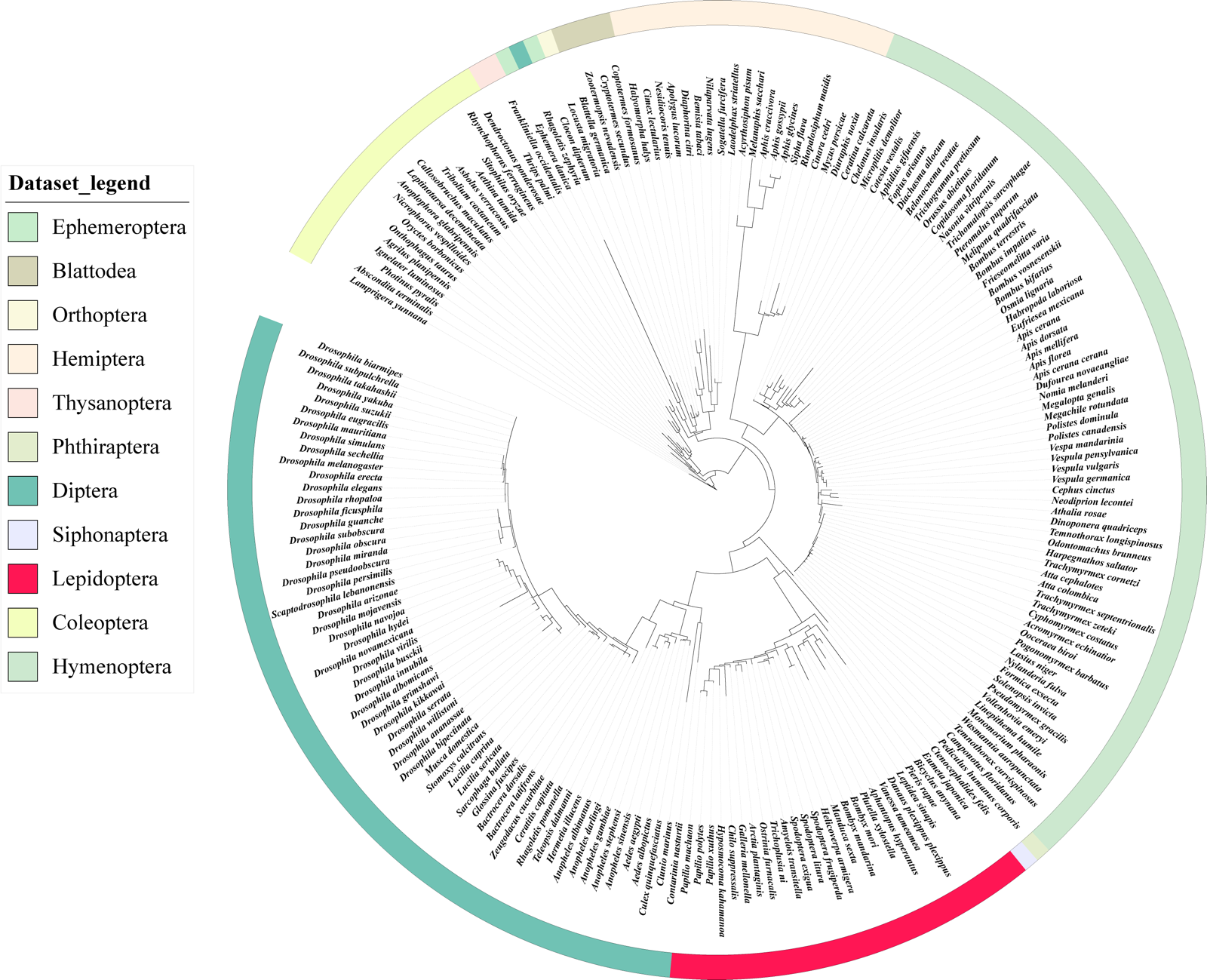

**Figure S3. Phylogenetic analysis of genes encoding METTL14**

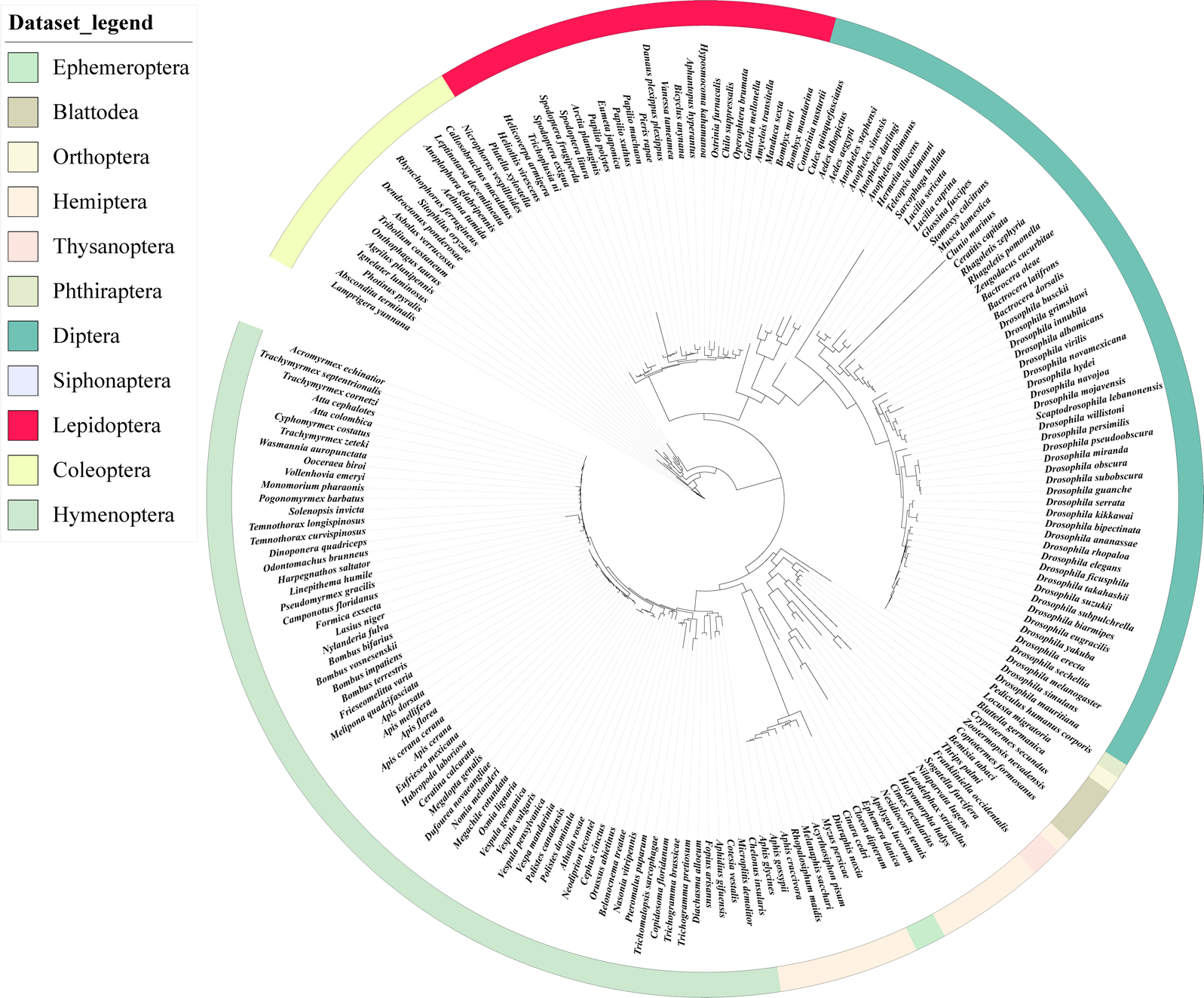

**Figure S4. Phylogenetic analysis of genes encoding FL2D**

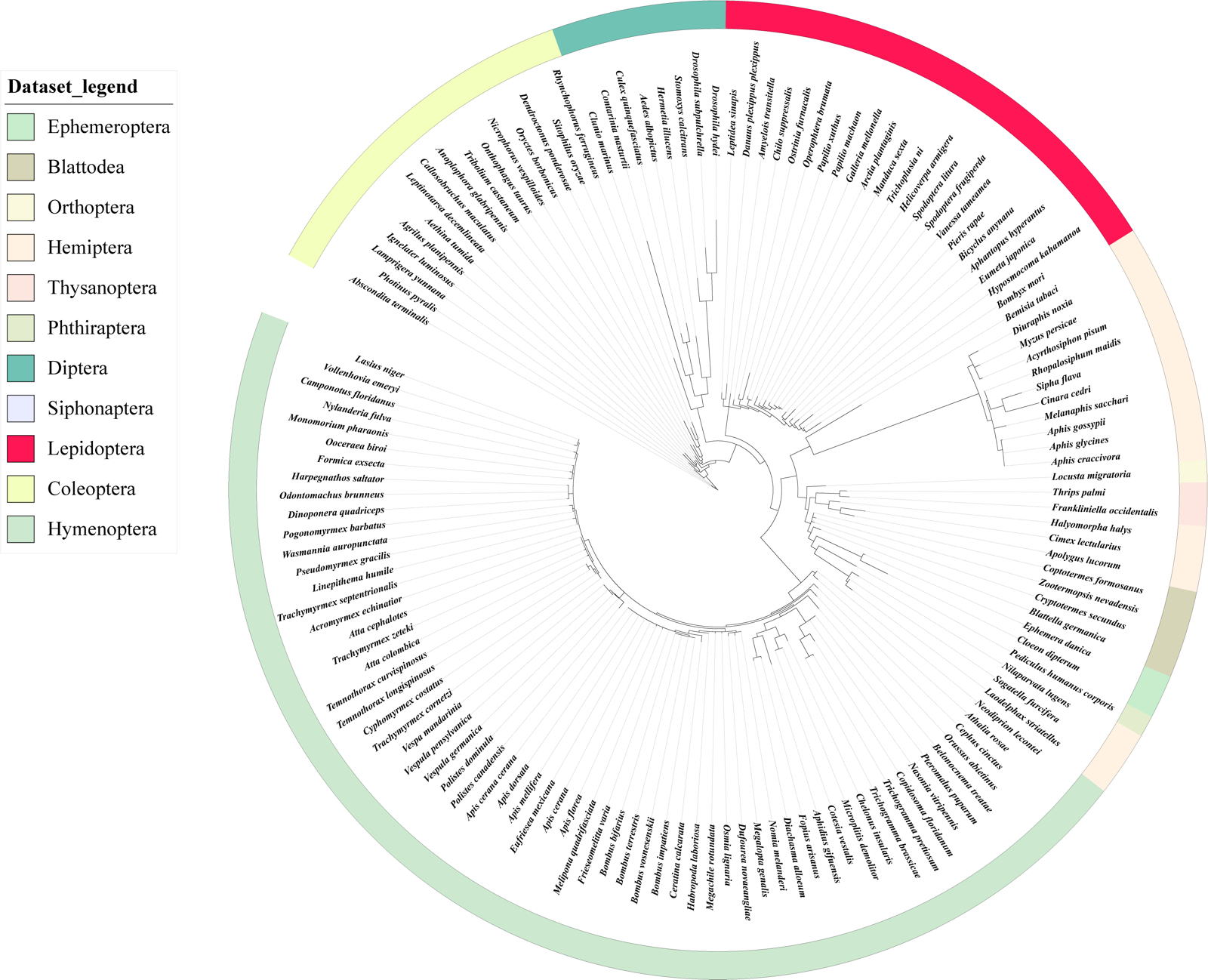

**Figure S5. Phylogenetic analysis of genes encoding YTHDF3**

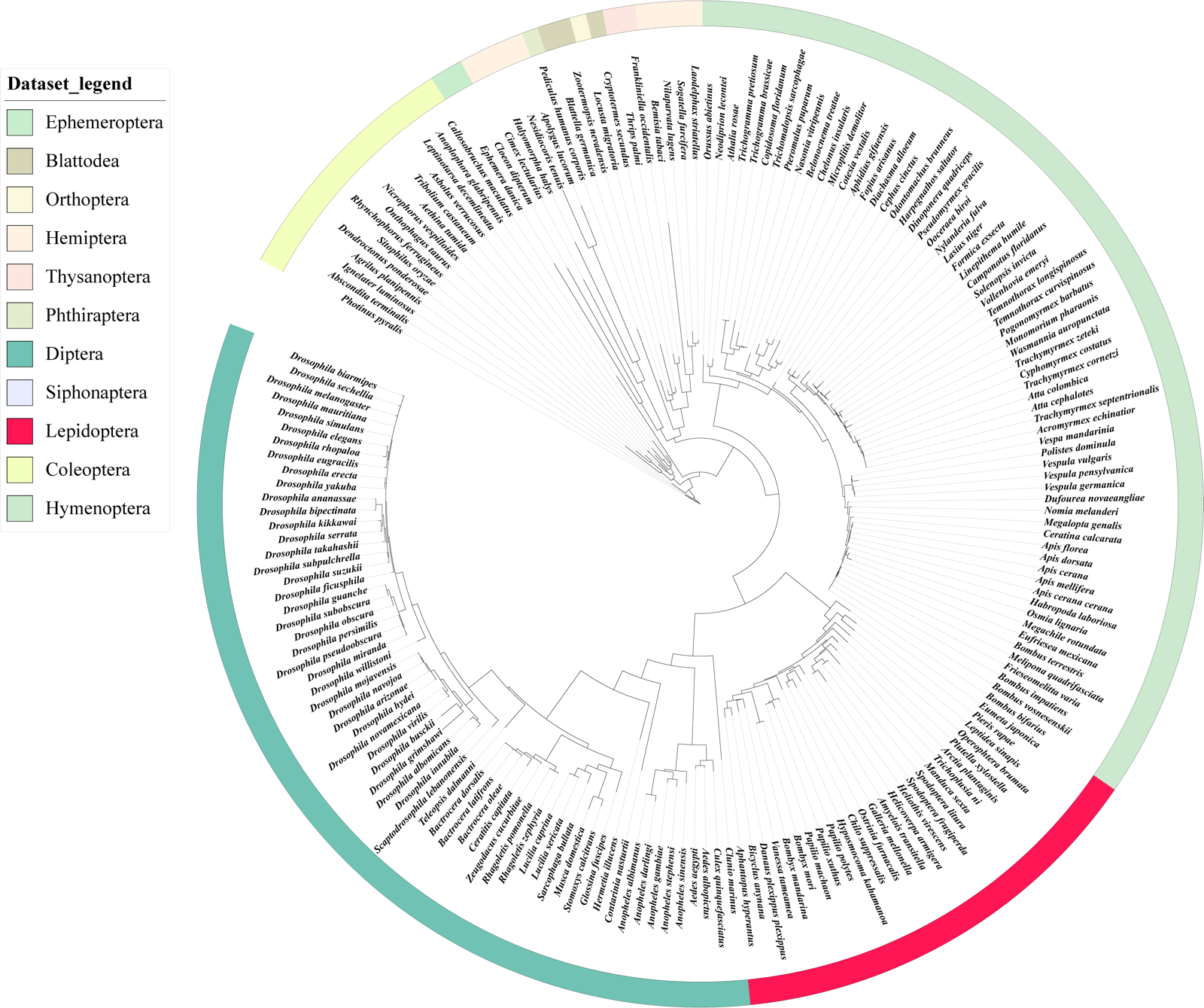

**Figure S6. Phylogenetic analysis of genes encoding YTHDC**

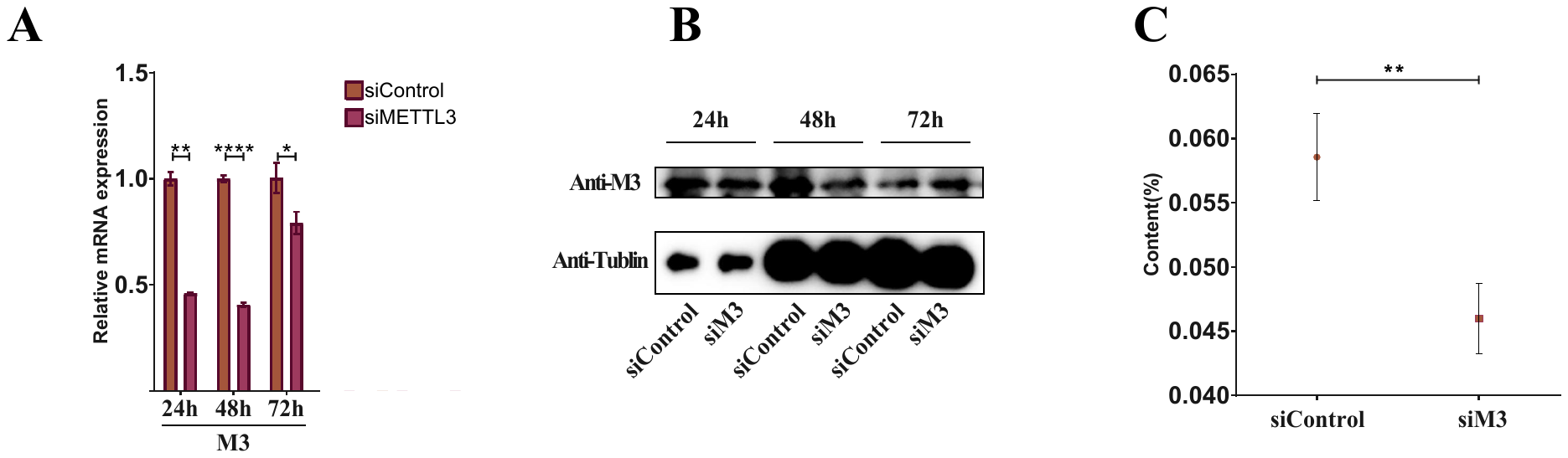

**Figure S7.** The m6A abundance of total RNA in siRNA injected eggs were determined at 48 h. The significance of difference was determined by Student’s t-test and denoted by *P < 0.05, **P < 0.01, and ***P < 0.001; n.s., not significant.

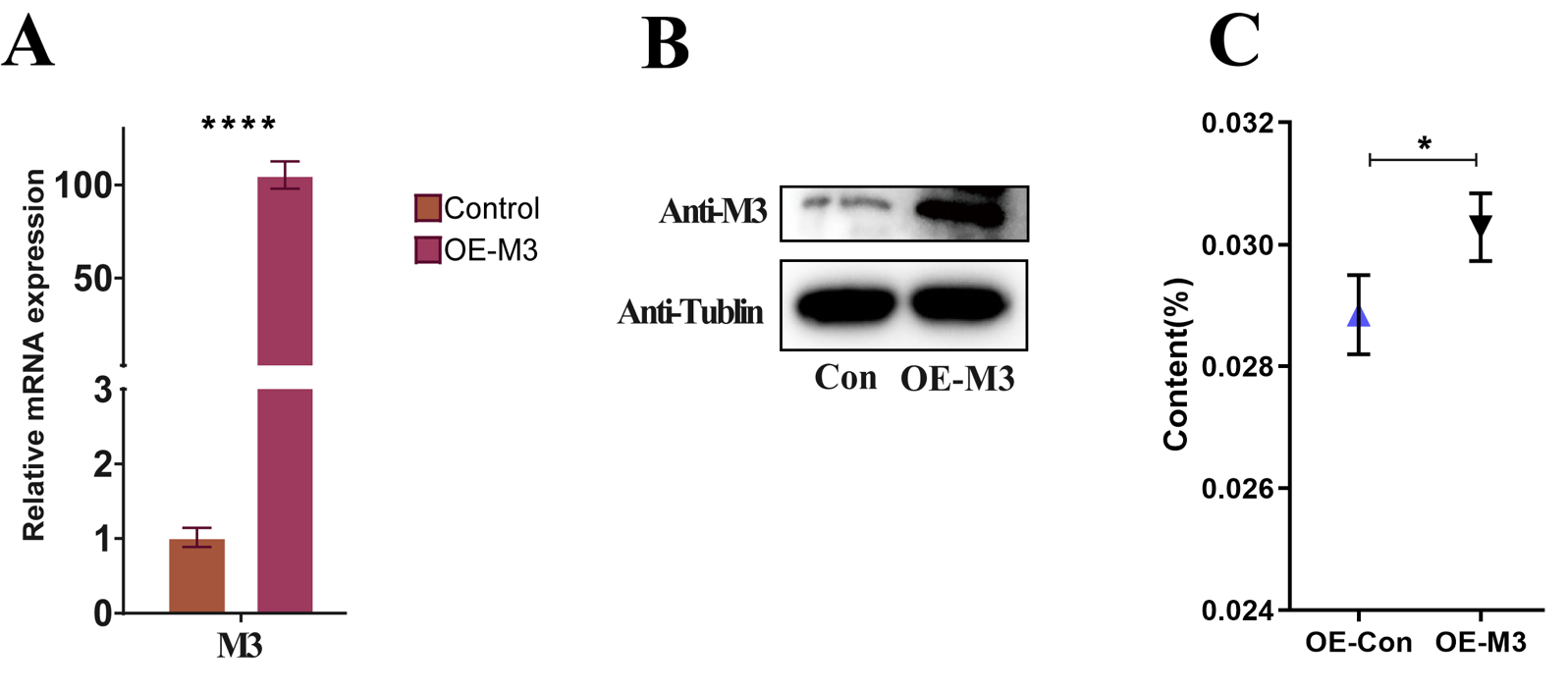

**Figure S8.** The m6A abundance of total RNA in BmN cells were determined. The significance of difference was determined by Student’s t-test and denoted by *P < 0.05, **P < 0.01, and ***P < 0.001; n.s., not significant.

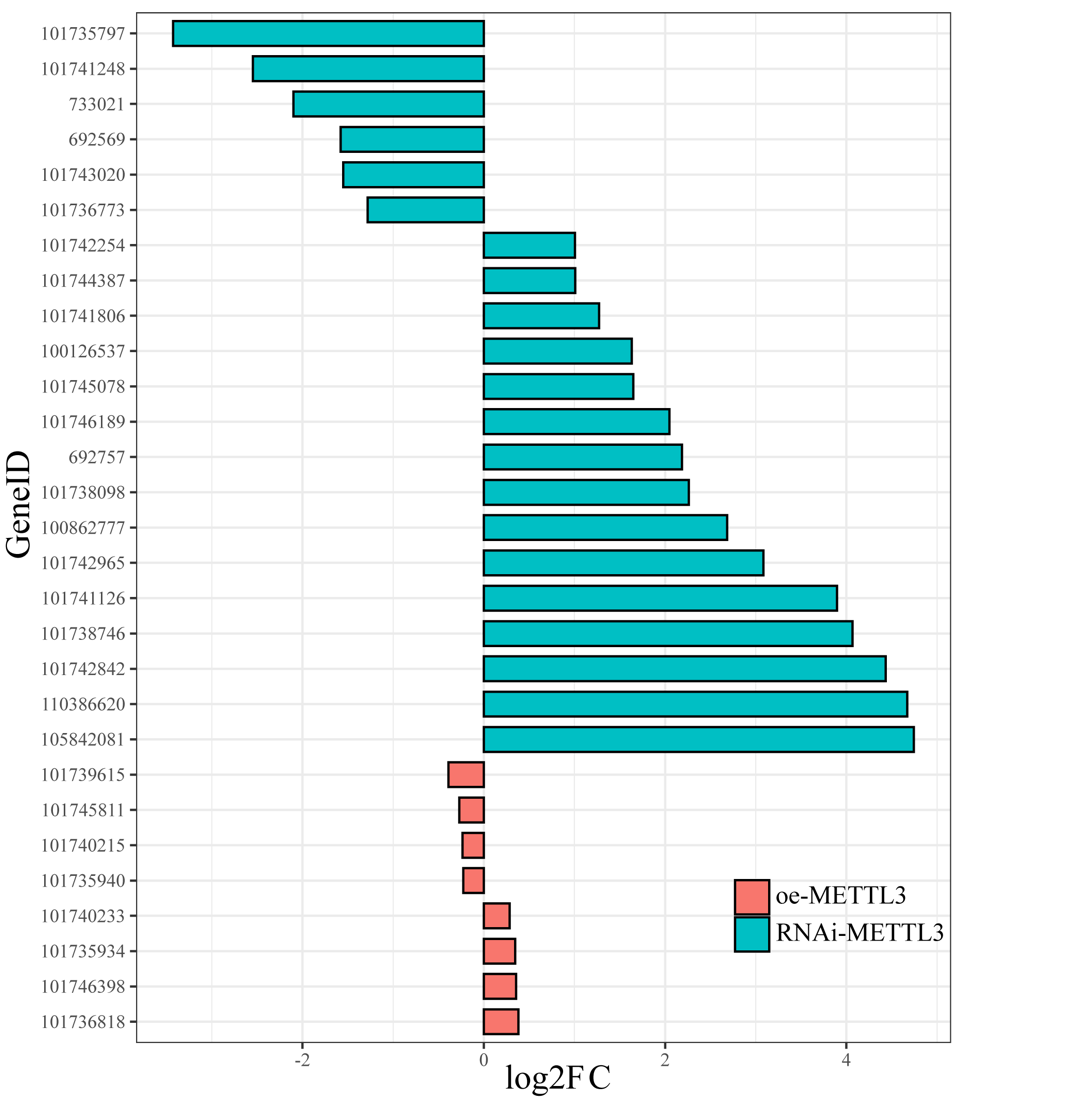

**Figure S9.** Analysis of differential genes (FDR<0.05) in pathways of Purine metabolism. Blue bar reprensent differential genes in RNAi-METTL3 in silkworm embryonic. Orange bar represent differential genes in OE-METTL3 in BmN cells.
